# Gene-environment interaction loci with signatures of mid- to late Holocene selection show evidence for diet-dependent genetic effects

**DOI:** 10.64898/2026.10.05.756704

**Authors:** Lotte Wijnhoven, Danat Yermakovich, Michael Dannemann

## Abstract

Environmental change is a major driver of adaptation, yet most evolutionary studies assume that genetic effects remain constant across environments. Recent biobank-scale studies have identified widespread gene-environment interactions (GxE), but their contribution to recent human evolution remains unclear. Here, we combined genome-wide GxE maps with allele-frequency trajectories spanning the last ~10,000 years to test whether environmentally responsive variants contributed to recent human adaptation. Among 45 independent loci, three showed significant signatures of directional selection. These loci, associated with urate, LDL-C and cholesterol levels, showed parallel frequency increases and overlapping acceleration points around 4,700-7,200 years ago. Selection-associated GxE loci were enriched for urate-related variants and for interactions involving meat and cheese consumption. Diet-associated variants showed a consistent interaction direction across loci. Notably, across candidate traits, GxE variants were enriched among significant GWAS loci with selection signatures. Furthermore, analyses across worldwide populations revealed broadly shared derived allele frequency increases and comparable temporal patterns. Taken together, our results suggest that gene-environment interactions present an important and previously underappreciated component of recent human evolution.

## Introduction

Since their emergence about 200,000 years ago, modern humans have encountered a variety of environmental conditions, including different climates, pathogens and shifts in diet that required them to adapt to those challenges^1^. Well-known examples of genetic adaptations to changing environments include the *LCT* locus, coding for lactase persistence, which has been closely linked to the shift to agriculture^2^, or malaria resistance associated with variants in the *DARC* gene, reflecting adaptation to pathogen exposure^3^. Historically, signatures of positive selection were inferred indirectly from genomic footprints using allele-frequency and haplotype-based approaches based on modern genomes^4^. In recent years, the increased availability of ancient DNA datasets has allowed for the reconstruction of empirical allele frequency trajectories, which provide a more direct framework to assess signatures of selection among variants associated with human phenotypes^5–9^.

The majority of modern human traits, however, are not determined by single variants, but are highly polygenic^10,11^. Thus, an evolutionary response to a changing environment often involves allele frequency changes at a variety of loci, something that is not generally taken into account during evolutionary analyses. Over the past decade, genome-wide association studies (GWAS) have enabled the study of the genetic basis of such complex traits, by identifying associations between a set of common genetic variants and a wide range of phenotypes within a cohort^12^. This has resulted in well-established sets of loci known to be associated with human diseases and non-disease traits. Even though GWAS have advanced our understanding of the genetic bases of complex traits and evolutionary forces acting on those traits, the methods used in such studies generally assume that genetic effects are fixed across environments. In recent years, however, an increasing number of studies have shifted the focus to investigate how environmental factors influence the genetic effects on human phenotypic variation through, so called, gene-environment interactions (GxE)^13,14^. As a result of several large-scale GxE studies, it is now well-established that these interactions are important for understanding complex traits and that they can systematically be detected within biobank-scale datasets^15–17^.

However, it remains unknown to what extent GxE variants have been targets of recent selection in humans and whether any traits or environments are particularly involved. To address this question, in this study, we combined genome-wide gene-environment interaction information from a recent biobank-based study^17^ with empirical allele frequency trajectories from the AGES database^18^ to assess whether signatures of directional selection among GxE loci reflect adaptation to environmental changes occurring over the past 10,000 years.

## Results

### Identification of GxE loci with signatures of selection

In this study, we aimed to explore the role of selection on genetic variants with signatures of gene-environment interactions (GxE). To address this task, we leveraged data from a recent study that mapped GxE on a genome-wide scale in UK Biobank (UKB) and Biobank Japan (BBJ)^17^. Namba and colleagues performed a screening of genetic variants with GxE signatures for a total of 47 traits, comprising 38 clinical biomarkers, that included anthropometric, metabolic, immune and blood measures as well as nine diseases that span cardiometabolic, inflammatory, renal and ocular disorders (Supplementary table 1). The GxE screening involved the testing of the contribution of nine environmental factors to the genetic effect sizes. These factors included age, sex, and lifestyle factors that involved beverage consumption, alcohol and food intake, smoking habits and physical activity (Supplementary table 2). The study identified 64 GxE across 26 traits and 45 distinct loci in UKB, and 36 interactions across 30 traits and 15 loci in BBJ, demonstrating that gene-environment interactions can be detected across a wide range of loci and traits.

We explored whether reported GxE loci show signatures of recent selection by screening GxE candidate variants in the AGES database^18^. Using genomes of ancient and modern humans of predominantly European ancestry, AGES provides reconstructed allele frequency trajectories over the past ~10,000 years and tests for the deviation of those trajectories from neutrality. For this analysis, we focused on GxE candidates identified in UK Biobank to match the predominantly European ancestry of the AGES cohort. We excluded four GxE variants that were located in the Major Histocompatibility Complex due to the complex linkage disequilibrium (LD) structure and pervasive selection signals in that region, which complicate the attribution of selection to specific loci or traits. Selection information for 51 of the remaining GxE variants was available in AGES. In addition, we included allele frequency data for four additional GxE variants that were not in the database but for which high-LD proxy variants were identified that were tested in AGES (UK Biobank r^2^> 0.95, Supplementary table 3, Methods). Hence, the combined set consisted of 55 candidate and proxy variants. Ten variant pairs showed elevated levels of LD (r^2^>0.5) and we collapsed each pair into one locus, resulting in 45 independent loci across the 55 candidate variants (Supplementary table 3).

**Table 1:**
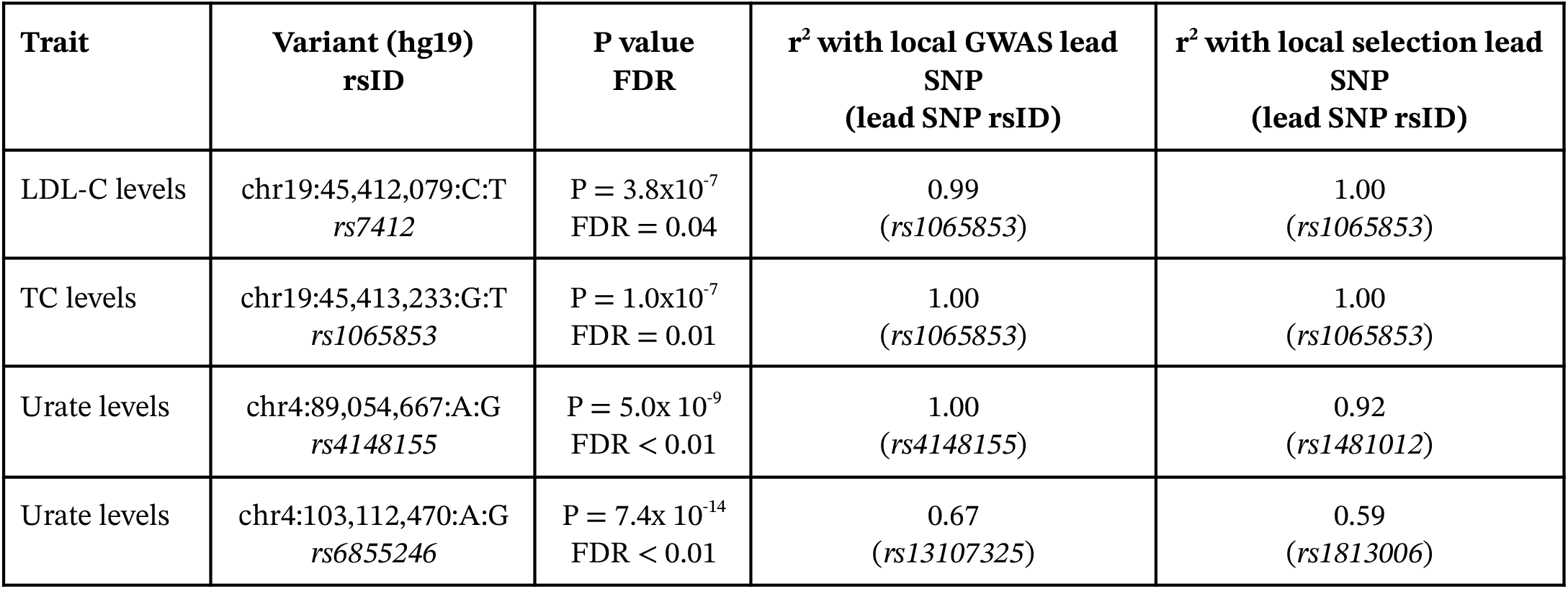
Overview of selection and GWAS statistics for GxE candidates. Shown are, for each of the candidate variants, the associated trait, the variant identifier (chromosome:position:reference:alternative), rsID, selection P value and FDR, the LD levels with both the GWAS lead SNP and the selection lead SNP.

| Trait | Variant (hg19)<br>rsID | P value<br>FDR | $r^2$ with local GWAS lead<br>SNP<br>(lead SNP rsID) | $r^2$ with local selection lead<br>SNP<br>(lead SNP rsID) |
| --- | --- | --- | --- | --- |
| LDL-C levels | chr19:45,412,079:C:T<br>*rs7412* | P = $3.8 \times 10^{-7}$<br>FDR = 0.04 | 0.99<br>(*rs1065853*) | 1.00<br>(*rs1065853*) |
| TC levels | chr19:45,413,233:G:T<br>*rs1065853* | P = $1.0 \times 10^{-7}$<br>FDR = 0.01 | 1.00<br>(*rs1065853*) | 1.00<br>(*rs1065853*) |
| Urate levels | chr4:89,054,667:A:G<br>*rs4148155* | P = $5.0 \times 10^{-9}$<br>FDR < 0.01 | 1.00<br>(*rs4148155*) | 0.92<br>(*rs1481012*) |
| Urate levels | chr4:103,112,470:A:G<br>*rs6855246* | P = $7.4 \times 10^{-14}$<br>FDR < 0.01 | 0.67<br>(*rs13107325*) | 0.59<br>(*rs1813006*) |

Four GxE variants spanning three independent loci showed significant allele frequency shifts over the past ~10,000 years (FDR<0.05, Table 1, Supplementary table 3, Methods). Two of the selection loci were located on chromosome 4 and associated with urate levels (*rs4148155*, hereafter Urate-locus1, *rs6855246*, Urate-locus2, Table 1). The third locus harbored two GxE variants, one associated with total cholesterol (TC) levels (*rs1065853*, TC-locus3) and the other with low-density lipoprotein cholesterol (LDL-C) levels (*rs7412*, LDL-locus3, Table 1). In all four instances, the derived allele is estimated at between 0.5 and 2% ~10,000 years ago and has increased to ~6-10% since (Figure 1; Supplementary figure 1). Three of the candidate variants showed a tight link to the local selection peak (all r^2^> 0.90 to the lead SNP). The fourth variant, associated with one of the urate-associated GxE candidates on chromosome 4, showed lower LD with the local selection lead SNP (r^2^= 0.59, Supplementary table 3).

**Figure 1:**
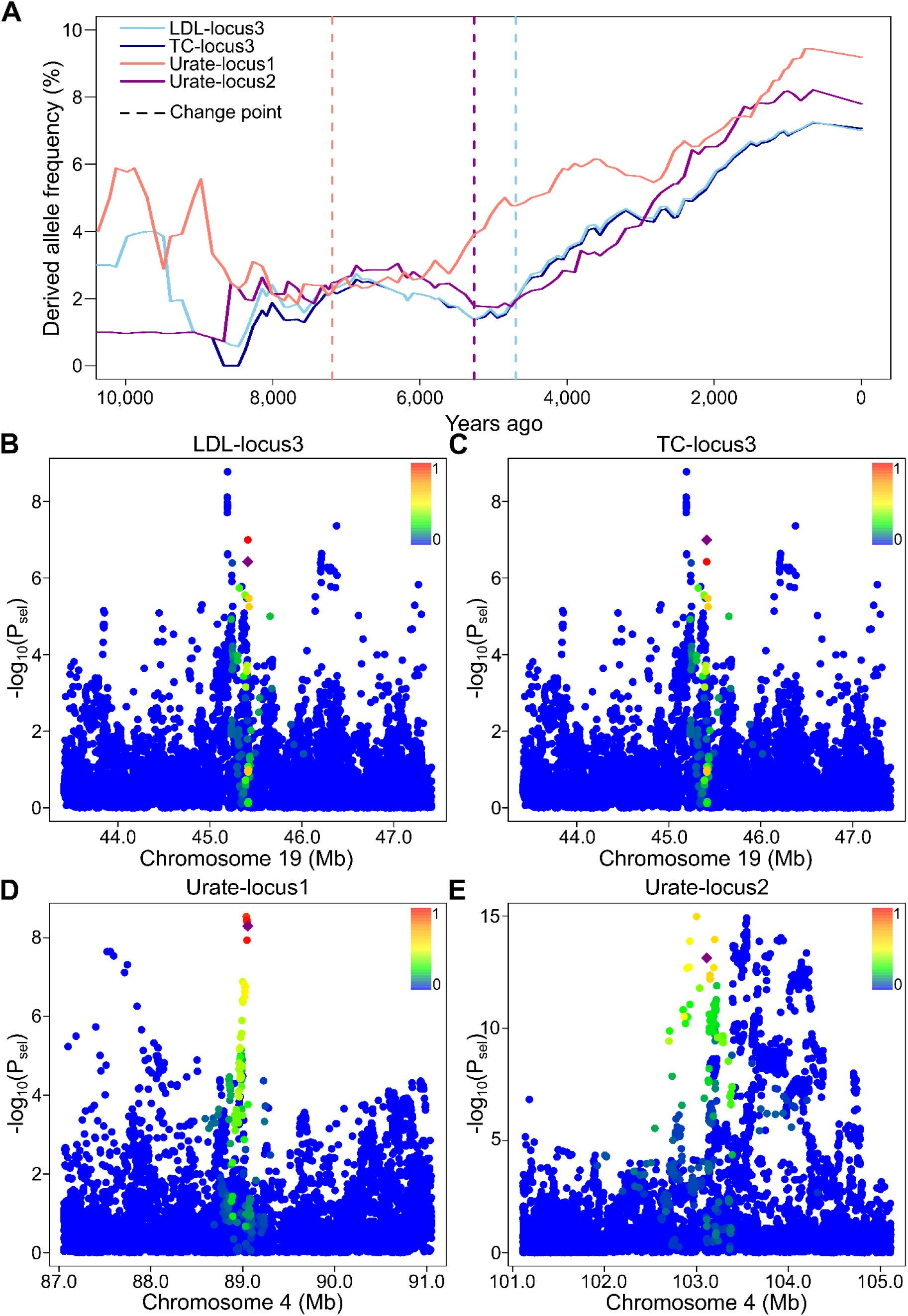
Selection signatures of four GxE variants with significant allele frequency changes over the last 10,000 years in the AGES database. **(A)** Empirical derived allele frequency trajectories for four GxE variants showing significant allele frequency changes over the last 10,000 years in the AGES dataset (solid lines, variant color scheme shown in the legend in the upper left of the panel). Dashed lines indicate the change points that partition each trajectory into two segments with the greatest difference in allele frequency change. LDL-locus3 and TC-locus3 share the same change point; consequently, their dashed lines overlap. **(B-E)** Manhattan plots of selection P values from the AGES cohort study (y-axis, −log10-transformed) for genomic regions (x-axis) encompassing each GxE candidate variant. Candidate variants are shown as purple diamonds, whereas all other variants are shown as circles and colored according to their r^2^ value with respect to the candidate variant (color scale shown in the upper right of each panel).

### Trait and environment enrichment for GxE candidate loci

Next, we explored whether selection-associated GxE loci showed enrichment for particular traits or environmental variables. Because several GxE variants are in LD and therefore not genetically independent, we performed enrichment analyses both at the variant level and after collapsing variants into independent loci. This distinction is important, as variant-based analyses may overrepresent signals from loci harboring multiple correlated variants, whereas locus-based analyses assume full redundancy within such regions. Consistent with this, environmental interaction profiles among GxE variants within the same locus showed moderate overlap (average pairwise Jaccard index=61.5%, Methods), a similarity that exceeded random expectation, as randomly generated sets showed overlaps of 15.3-44.2% (95% confidence interval, permutation test, P=1.8×10^−4^, Methods), indicating that environmental interaction signals are only partially shared across LD proxies and not fully captured at the locus level. We therefore decided to interpret the variant- and locus-based enrichment results as providing upper and lower bounds, respectively, with the true enrichment signal likely lying between both estimates.

In total, Namba et al report four GxE loci that were associated with urate levels. The observation that two of the three selection loci were linked to two of these four urate-associated loci was unexpectedly high, at both the variant (OR=20.9, P=0.023) and locus level (OR=31.5, P=0.018, Figure 2a,b). This result was in contrast to TC and LDL-C–associated loci, which did not show an enrichment for selection signatures (TC, variant-level: P=0.38 / locus-level: P=0.36; LDL-C, variant-level: P =0.27 / locus-level: P=0.25, Figure 2b, Supplementary figure 2).

**Figure 2:**
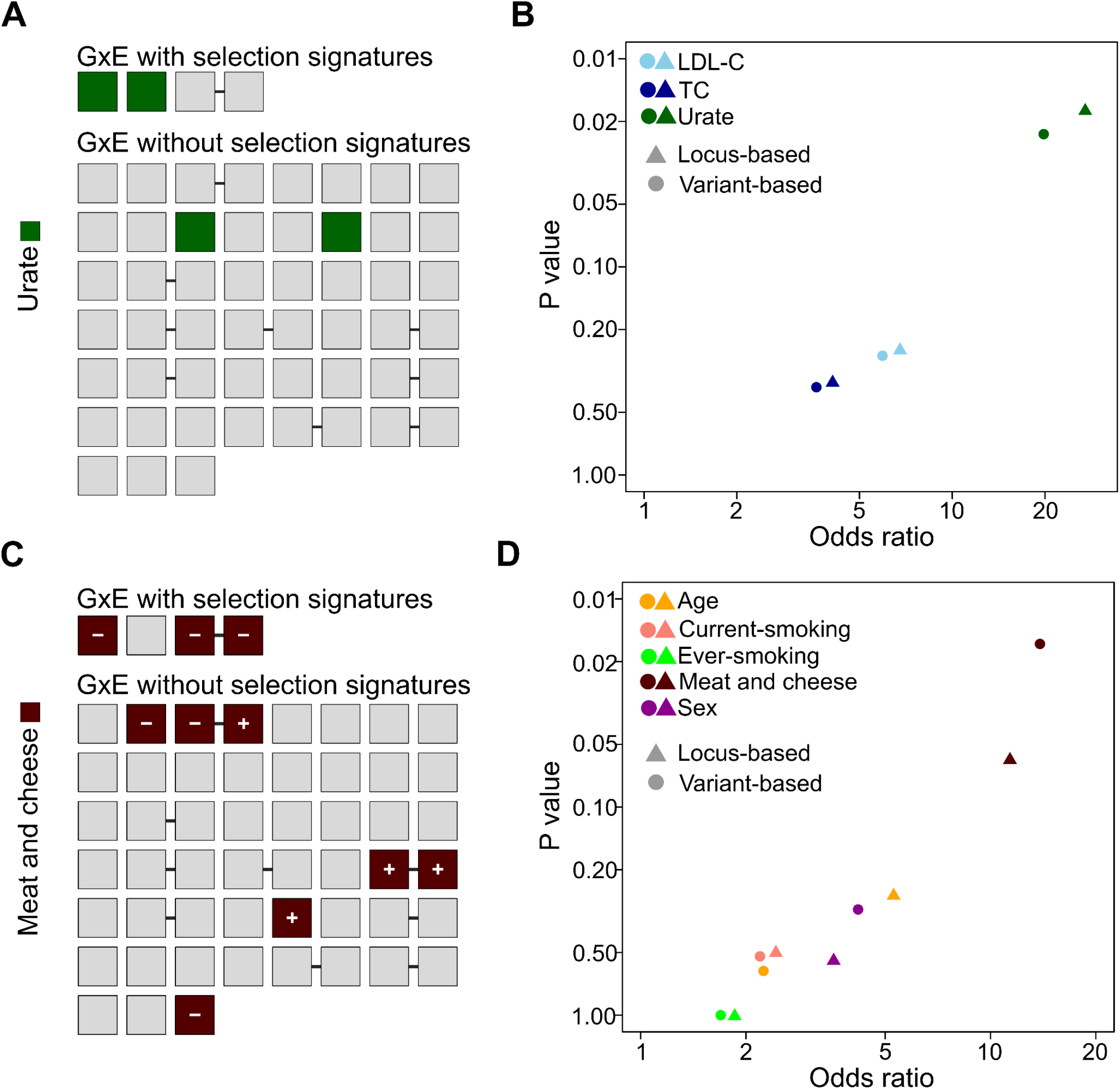
Trait and environment enrichment of GxE variants with signatures of selection. **(A,C)** GxE variants are shown as squares and partitioned into two sets: one comprising all four candidate variants (upper set) and a second comprising all 51 variants without signatures of selection (lower set). Variants within both sets are ordered according to their genomic position, and variant pairs belonging to the same locus are connected by a black line. Variants associated with urate **(A)** and meat and cheese diet **(C)** are highlighted in green and brown, respectively. In addition, meat and cheese diet-associated variants are marked with a ‘+’ or ‘-’ symbol to indicate the sign of the interaction beta with respect to the derived allele effect. **(B,D)** Volcano plots showing Fisher’s exact test enrichment P values (y-axis, log-scale) and odds ratios calculated using the Haldane-Anscombe correction (x-axis, log-scale) for all traits **(B)** and environments **(D)** associated with at least one of the four candidate GxE variants. Color and shape schemes for traits, environments, and test mode (variant-based, locus-based) are shown in the upper left of each panel.

Furthermore, Namba et al tested gene-environment interactions for nine environmental variables and reported for each GxE variant a subset of contributing variables. For a given GxE candidate variant, environmental variables were evaluated in a stepwise approach for their contribution to the interaction signature. The four GxE variants mapping to the three selection-associated loci were collectively associated with five of these nine contributing environmental variables, each observed in at least one selection-associated variant. Those contributing variables were: age (3 associated selection variants across 2 selection loci), current (1 variant) and ever smoking status (1 variant), meat and cheese diet (3 variants across 2 loci) and sex (4 variants across 3 loci). Given that only 10 of the 55 GxE variants with available AGES information included a meat and cheese diet interaction, the observation that three of these variants showed signatures of selection was unexpectedly high (OR=17.3, P=0.016, Figure 2c,d). This result was still borderline significant when we performed the same calculation based on independent loci (OR=13.3, P=0.059). None of the other four variables showed an enrichment among selection-associated GxE variants and loci (all P>0.25, Supplementary figure 2, Supplementary table 4).

Interestingly, all three interactions involving selection-associated GxE variants and meat and cheese diet showed negative interaction effects, when polarizing the genetic and environmental effect to the derived alleles, indicating a consistent direction of environmental modulation across loci. Three of the seven non-selection GxE candidates interacting with meat and cheese showed this direction with respect to the derived allele, whereas the remaining four showed interaction effects in the opposite direction. None of the other two environmental variables repeatedly associated with GxE selection candidate variants (age and sex) showed a consistent directional effect. Notably, observing at least three variants or two loci associated with meat and cheese and all showing a consistent directional effect is unlikely under random expectations (variant-based P=0.037, locus-based P=0.017, permutation test, Methods, Supplementary figure 2c).

### GWAS signatures of GxE candidate loci

Next, we explored the traits associated with selection-linked GxE loci in terms of their overall polygenicity and signatures of selection, and assessed how the GxE selection variants fit within this broader trait architecture. First, we assessed overall polygenicity by identifying independent significant association loci for each of the three traits associated with GxE selection variants, using publicly available UK Biobank GWAS summary statistics (Methods).

All three traits showed elevated levels of polygenicity, with 212 (TC), 192 (LDL-C), and 314 (urate levels) independent significant loci, respectively. We found that all four selection-associated GxE variants were linked to one of those genome-wide significant association loci of their respective traits (Supplementary table 3). Overall, 12 of the 14 GxE variants associated with urate, LDL-C, or TC reported by Namba et al showed genome-wide significant associations in the corresponding trait GWAS, suggesting this feature being highly prevalent among GxE variants (Supplementary table 3). Furthermore, three of the four GxE selection variants were in high LD with the local GWAS lead variant (r^2^> 0.99), while the fourth variant also showed moderate LD (r^2^= 0.67), placing all four variants in close proximity to the local association signal (Table 1).

We found that among significant GWAS loci for LDL-C, TC and urate, between 4 and 7% harbored variants with signatures of selection (including all variants with r^2^> 0.5 to the locus lead SNP and selection FDR<0.05). This proportion is significantly lower than the one observed for GxE variants across the three traits (36.4%, P=6.0×10^−4^, OR=8.4, Mantel-Haenszel test, Methods), suggesting an enrichment of selection signals among GxE variants relative to the broader trait architecture. When considering traits individually, this enrichment was primarily driven by urate (P=6.2×10^−3^, OR=43.7, Fisher’s exact test), whereas TC and LDL-C did not show significant enrichment on their own (OR>1, P>0.20).

Next, we explored whether the level of polygenicity and the proportion of selection-associated GWAS loci differed systematically compared to other traits tested by Namba et al, potentially favoring candidate traits and biasing our inferences. To this end, we quantified the number of significant loci and the proportion showing signatures of selection for 31 additional traits for which GWAS summary statistics were available and that were analyzed by Namba et al. We found that while neither of the three candidate traits were an outlier in terms of polygenicity nor in the absolute number of selection loci, both LDL-C and TC showed the second and third highest proportions of selection loci among all tested traits (Supplementary figure 4).

### Evolutionary inferences of GxE candidate loci across populations

To better understand the implications of our inferences for present-day populations of primarily non-European ancestry, we explored the frequency distribution of the four GxE candidate variants across global populations. In European populations of the 1,000 Genomes cohort, these variants were observed at frequencies between 1 and 14% (Supplementary table 5, Figure 3a-c, Methods). With the exception of the urate-associated variant Urate-locus2, which was not detected in three East Asian populations, the GxE selection candidate variants segregated across all five continental super-populations (Africa, the Americas, East Asia, Europe, and South Asia), in some cases reaching frequencies of up to 31% (Supplementary table 5, Figure 3a-c). Despite their presence across populations, none of the candidate variants are among the six GxE loci that were found to overlap between the UK Biobank and Biobank Japan by Namba et al. However, we note that the environmental variables partly differed between the GxE analyses in the two biobanks, which may have contributed to the lack of replication of some candidate loci across populations.

**Figure 3:**
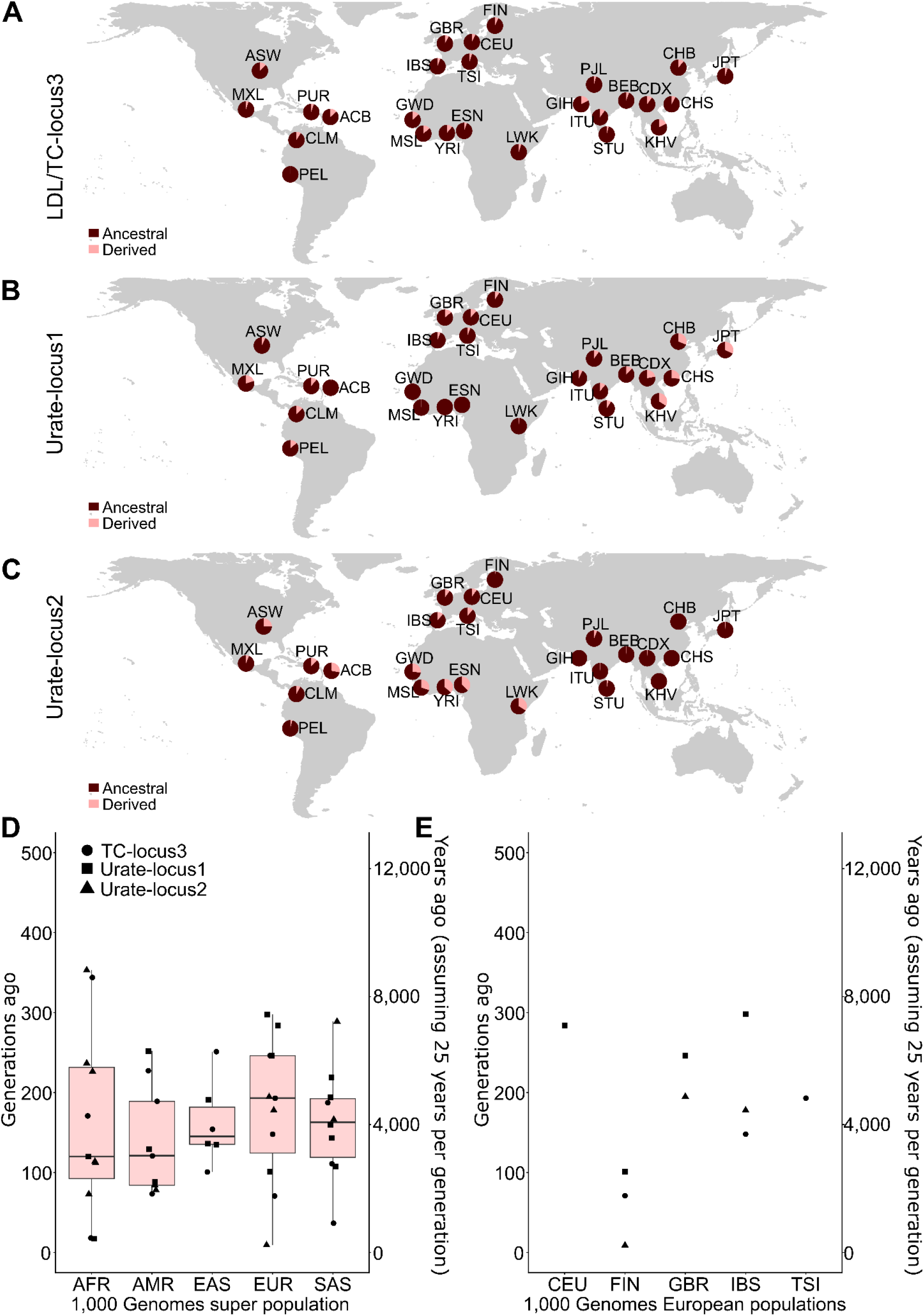
Cross-population genomic inferences for candidate GxE variants. **(A-C)** Derived allele frequencies for GxE candidate loci across 26 populations from the 1,000 Genomes Project are displayed (orange slice of each pie chart). For LDL/TC-locus3 **(A)**, allele frequencies were calculated as the average frequency of the two associated variants within each population. **(D,E)** Estimates of change points in allele frequency trajectories reconstructed using CLUES are shown for three testable GxE variants across five superpopulations **(D)** and five European populations **(E)** from the 1,000 Genomes Project. Only significant change points partitioning trajectories into two segments with significantly different slopes (P<0.05) are shown. Panel D additionally includes boxplots summarizing the distribution of change-point estimates across superpopulations, whereas panel E shows only population-specific estimates. The y-axes are shown in generations (left axis) and years (right axis), assuming 25 years per generation.

Reconstructed allele frequencies not only provide information about selection signatures but also offer a powerful means to narrow down the onset of selection episodes. We therefore performed a breakpoint detection test, based on a piecewise linear modeling framework, to identify time periods during which trajectories showed significant shifts in slope (Methods). This test assesses whether partitioning each allele frequency trajectory into two segments results in a significantly better fit of the data than a single-line model. We found that such shifts in slopes of the allele frequency trajectories for all four candidate variants occurred between approximately 4,700 and 7,200 years ago, after which the allele frequencies increased more rapidly (P<0.001, Methods, Figure 1a).

Next, we explored whether any of our candidate variants showed comparable derived allele frequency changes in other world-wide populations, potentially reflecting shared selection signatures. In the absence of ancient genomic data from non-European populations at a scale sufficient to robustly reconstruct empirical allele frequency trajectories comparable to AGES, we employed an alternative approach that does not rely on ancient DNA. This approach is based on the CLUES algorithm, which leverages local genealogical trees to infer past allele frequency trajectories and test for signatures of selection. We generated local trees for all 26 populations of the 1,000 Genomes dataset using Relate and applied CLUES to our candidate variants (Methods). As LDL-locus3 did not pass the Relate quality filters, it was excluded, and the analysis was performed on three of the four GxE variants. Across 65 of 66 tested variant-population pairs, we observed a consistent increase in the frequency of the derived allele over the past 500 generations, which are equivalent to ~12,500 years when assuming a generation time of 25 years. Next, we applied the same breakpoint identification method as for the AGES data and found that 47 of the 66 allele frequency trajectories showed a significant breakpoint (P<0.05, permutation test, Supplementary figure 3, Supplementary table 8, Methods). A total of 25 of 27 trajectories in Eurasian populations showed a steeper allele frequency increase following the breakpoint (P=5.65×10^−8^, binomial test, Supplementary figure 3), whereas in African and American populations, the proportions of trajectories with a steeper increase before the breakpoint (55%) and after the breakpoint (45%) were roughly equal (P=0.82, binomial test). These results suggest that accelerations in allele frequency change in Eurasia are preferentially shifted toward more recent time periods, while such temporal structuring is less pronounced in other populations. Overall, for European populations, the median breakpoint occurred 192 generations ago, corresponding to approximately 4,825 years BP assuming a generation time of 25 years - an estimate close to the values observed in the AGES database. Furthermore, within European populations, we observed a notable difference in breakpoint estimates between the Finnish population, which ranged from 9 to 101 generations ago (225 to 2,525 years BP), and other European populations, which ranged from 148 to 298 generations ago (3,700 to 7,450 years BP). Overall, breakpoint estimates for European populations were comparable to those of non-European populations (Wilcoxon rank test, P=0.35), which showed a median breakpoint of 139.5 generations ago (3,487 years BP) and ranged from 17 to 353 generations ago (425 to 8,825 years BP) (Figure 3d,e).

### Phenotypic consequences of GxE variants

Next, to assess the broader phenotypic implications of GxE variants with selection signatures, we examined their presence across GWAS. We found that associations spanned a wide range of metabolic traits (Supplementary table 6). However, despite shared environmental responsiveness and evidence of selection, variants differed substantially in their degree of pleiotropy and trait specificity.

The allele that increased in frequency for Urate-locus1, which mapped to the lead urate GWAS locus in UK Biobank (Supplementary table 3), was associated with higher urate levels. The variant is located near *ABCG2*, a well-established urate transporter implicated in hyperuricemia, gout, and kidney stone risk, and has previously been associated with multiple urate- and kidney-related phenotypes. Compared to the other selected GxE loci, the locus showed relatively limited pleiotropy, with secondary associations primarily involving kidney function, metabolic traits, and inflammatory markers. Interestingly, Urate-locus1 was among the selected GxE variants showing interaction with meat and cheese intake, where the genetic effect was stronger in individuals with a lower meat and cheese intake. Although the evolutionary implications remain uncertain, this pattern is compatible with environment-dependent modulation of disease-associated urate effects. While Urate-locus1 lies within an intron of *ABCG2*, it is in perfect LD (r^2^=1) with *rs2231142*, a missense variant in *ABCG2*, that may be a more likely functional driver of the observed phenotypic effects (Supplementary table 7).

The second urate-associated GxE candidate, Urate-locus2, showed several markedly different features compared to Urate-locus1. First, the allele associated with increasing frequency over time was linked to lower urate levels, opposite to the pattern observed at the *ABCG2* locus. Second, Urate-locus2 exhibited substantially higher levels of pleiotropy, with associations spanning musculoskeletal, cardiometabolic, and protein/blood chemistry traits, as well as weaker associations involving immune-related phenotypes, blood cell traits, lipid composition, and brain traits (Supplementary table 6). The variant is located in proximity to *SLC39A8*, a gene with broad physiological functions related to metal ion transport and systemic metabolism. Urate-locus2 was the only one of the four selection-associated GxE variants that did not show interaction with meat and cheese intake, suggesting heterogeneity in the environmental contexts linked to selection across urate-associated loci. And while Urate-locus2 was annotated as intergenic, it showed elevated levels of LD (r^2^=0.66) with *rs13107325*, a missense variant for *SLC39A8* and a potentially more likely functional candidate for the associations observed at this locus (Supplementary table 7). Together, these observations are consistent with a more highly pleiotropic and systems-level form of metabolic adaptation, potentially involving trade-offs across multiple physiological pathways rather than a more trait-focused evolutionary effect as observed for Urate-locus1.

Both selection-associated GxE variants on chromosome 19 (LDL-locus3 and TC-locus3) mapped to the *APOE* locus, a central regulator of lipid transport and metabolism. The frequency-increasing alleles of both variants were associated with lower LDL-C and TC levels, respectively, and both showed gene-environment interactions involving meat and cheese intake. Both *APOE*-associated variants showed broad pleiotropy, with trait associations primarily centered on lipid biology and cardiovascular phenotypes, but also extending to body composition, blood traits, inflammatory and protein biomarkers, neurodegenerative diseases such as Alzheimer’s, as well as medication- and diet-related traits (Supplementary table 6). Both variants are in high LD (r^2^=0.99), and as LDL-locus3 is a missense variant in *APOE*, it is likely the more plausible functional candidate, with both variants tagging the same underlying signal (Supplementary table 7). Together, these observations are consistent with a more general systems-level role of the *APOE* locus in controlling energy allocation, lipid processing, and metabolic homeostasis, potentially involving context-dependent trade-offs between early-life metabolic benefits and later-life disease risks.

## Discussion

In this study, we investigated evidence for selection acting on variants with gene-environment interaction signatures. Although recent large-scale biobank data enable genome-wide identification of GxE effects, the number of detectable loci remains limited. Within the constrained set of a recent study, we identified four variants across three independent loci showing signatures of recent selection, exemplifying that some environmentally responsive variants have been shaped by recent evolutionary pressures. Two of these variants were linked to variation in urate levels, whereas the remaining two, that mapped to the same locus, were associated with LDL-C and TC levels.

Despite differences in genomic location and associated traits, the four selection-associated variants share several striking characteristics. One of the strongest similarities is their allele frequency trajectories in the AGES cohort. All four variants were segregating at low frequencies of less than 2% around 10,000 years ago and have since shown comparable increases to levels of around 6-9%. Even more striking in this context is their shared burst in allele frequency at a similar time point between approximately 4,700 and 7,200 years ago.

Furthermore, the four candidate variants were collectively associated with five environmental variables. Among these, only one - meat and cheese intake - showed evidence of enrichment for GxE with selection signatures. Notably, the three variants linked to this variable showed a consistent interaction direction, with genetic effects shifting in the negative direction with increasing meat and cheese intake. Finally, consistent with the majority (80%) of GxE variants reported by Namba et al, all four candidate variants were genome-wide significant hits in their respective trait GWAS.

Nevertheless, despite their shared association with urate levels, the two urate-associated variants differed markedly in both effect direction and degree of pleiotropy. While Urate-locus1 mapped to the top-associated locus and was linked to higher urate levels, Urate-locus2 showed a weaker association signal and was associated with lower urate levels. In addition, Urate-locus1 exhibited relatively limited pleiotropy, with secondary associations primarily involving gout and kidney stones, whereas Urate-locus2 showed a much broader range of pleiotropic associations across multiple biological systems. Notably, Urate-locus2 was also the only candidate variant not associated with meat and cheese intake as an environmental factor. At the same time, urate stood out more broadly across analyses, showing enrichment both among selection-associated GxE loci and among GWAS variants with signatures of selection, whereas no comparable enrichment was observed for TC or LDL-C.

Metabolic traits have long been recognized as targets of selection in recent human evolution^19,20^. In particular, dietary changes have frequently been invoked as a driving selective pressure, for example in the context of lactase persistence^21^, fatty acid metabolism^22^, and broader hypotheses of metabolic adaptation such as the “thrifty genotype” model^23^. However, direct evidence for signatures of selection among complex trait-associated variants with environmental interactions has remained limited.

In our study, we provide evidence for such a link. Across our candidate loci, we observe consistent signatures of selection together with environment-dependent effects that are particularly strongly linked to dietary factors. The enrichment of meat and cheese intake among environmental variables, in combination with the directional effects associated with our candidate loci, suggests that the fitness consequences of these variants may have depended on dietary exposure. Previous studies highlighting metabolism as a target of selection in humans have often proposed dietary change as a driving force^19^. Gene-environment interactions, as investigated in this study, may represent an additional layer through which such evolutionary processes can be studied.

To characterize these selection signatures, we analysed allele frequency trajectories over the past ~10,000 years. The trajectories of our candidate variants were not only strikingly similar, but also allowed us to narrow down the potential onset of selection, with accelerated allele frequency increases occurring between approximately 4,700 and 7,200 years ago. This time period in human history has been characterized by several cultural changes, including a transition towards pastoralism, which involved a shift in diet towards increased consumption of animal-derived products as well as greater exposure to zoonotic pathogens^24,25^. In addition, human populations became more sedentary and increasingly lived in higher-density settlements^26^. These populations were also characterized by increased Steppe-related ancestry due to large-scale demographic movements^27^. Together, these factors represent substantial changes in environmental conditions, which likely altered the landscape of genetic effects, particularly for traits sensitive to these transitions^6^. Our results are consistent with such shifts, where changes in diet may have modified genetic effects and, consequently, their fitness consequences. Overall, the shared allele frequency changes, the similarity in their timing, and their distribution across distinct genomic loci suggest that they may reflect a response to a common selective pressure. However, the heterogeneity across loci in terms of their phenotypic links implies that, while coordinated, this signature more likely represents a complex adaptive response rather than classical polygenic selection.

This complexity is particularly evident for the two urate-associated loci, which differ in their levels of pleiotropy, the direction of their association with urate levels, and the diversity of their environmental interactions. More generally, urate emerged as an intriguing trait in our study. In addition to the locus-specific differences described above, urate-associated GxE variants were enriched among both GxE selection loci and selection-associated GWAS loci, highlighting that this trait stands out more broadly across analyses. Urate levels are strongly linked to diet, as they reflect the processing and breakdown of purines, which are abundant in many foods, including a range of animal-derived products^22^. Urate levels are substantially higher in humans and other great apes due to the loss of uricase activity following pseudogenization of the UOX gene approximately 15-20 million years ago^28^. These elevated levels have been proposed to promote fat accumulation and enhance energy storage, which may have been advantageous during periods of food scarcity^29,30^. They may also contribute to maintaining blood pressure under low-salt conditions, such as during dietary transitions, heat stress, or dehydration^31^ and serve as an important antioxidant^32^. However, elevated urate levels are also associated with well-known disease trade-offs, including increased risk of gout and kidney stones^33^. Together, these features suggest that urate represents a plausible target in the context of our study, as diet-dependent environmental changes may have altered genetic effects on urate levels, with potential fitness consequences that could have been acted upon by natural selection.

Furthermore, previous studies of response eQTLs have shown that genetic effects on gene expression can be dependent on the pathogenic landscape, and that loci exhibiting such signatures are often associated with signals of selection or admixture^34–37^. Our results align closely with this work and extend the concept beyond molecular phenotypes, suggesting that environment-dependent genetic effects may also contribute to recent adaptation at the level of complex traits.

While these studies underscore the importance of environment-dependent genetic effects, their identification remains challenging. For example, the analysis of variance quantitative trait loci (vQTLs) has emerged as an important approach to identify footprints of GxE variants in recent years^38,39^. At the same time, vQTLs have been shown to miss a substantial fraction of GxE loci^17^ and recent work has suggested that they are associated with a considerable proportion of statistical artifacts^40^. In contrast, our approach uses variants with explicitly modelled gene-environment interactions and therefore provides a more specific framework to assess such interactions. In addition, our locus definition creates some ambiguity in the enrichment analysis, due to linkage disequilibrium between variants within a locus that partially but not fully overlap in their associated environmental variables. Nevertheless, the general overlap of results based on variant- and locus-level analyses suggests that our results are robust to this condition.

Furthermore, our analyses were based on Western Eurasian populations with a large European ancestry component, with GxE information derived from the UK Biobank and selection inferences from the AGES cohort. While this introduces a bias in the detection of candidate variants, several lines of evidence suggest that key aspects of our findings can be extrapolated to other populations. First, the four selection-associated GxE variants were detected in nearly all tested global populations. In addition, our CLUES-based analyses revealed a consistent increase in derived allele frequencies for three GxE variants across 65 of 66 variant–population pairs over the past ~500 generations, indicating broadly shared directional frequency changes across diverse populations. More importantly, a substantial fraction of these trajectories showed evidence of breakpoints that mark an acceleration in allele frequency change, with timing estimates in many populations aligning closely with those observed for the GxE candidate variants in the AGES dataset. This overlap of breakpoint estimates across multiple populations and with independent ancient DNA-based inferences supports the presence of a shared phase of accelerated allele frequency change. At the same time, we observed structured differences in the temporal dynamics of these signals. Notably, breakpoints in Eurasian populations were predominantly associated with more recent accelerations in allele frequency change, whereas African and American populations showed a less pronounced temporal structure. This may reflect differences in the timing and nature of major subsistence transitions across regions, such as the independent onset and intensification of agriculture^41,42^. Further support for this interpretation is provided by the delayed breakpoint estimates observed in the Finnish population, which are consistent with its later and more gradual transition to agriculture and pastoral subsistence compared to other European populations^43^. Together, these results suggest that while the underlying selective pressures on GxE candidates may be widely shared across populations, their temporal dynamics are consistent with differences in the timing and nature of agricultural and dietary transitions across regions.

Finally, we would like to acknowledge the limited number of GxE variants with selection signatures. This observation is partially due to the challenges to robustly detect variants that show gene-environment interactions in the first place^44^. However, despite the small number, we found that GxE variants that also show selection signals do not appear to represent random subsets. Instead, they show strong associations with traits and environments that are consistent with recent selection, such as metabolic traits and dietary exposures. Hence, GxE loci, while being rare, might represent a particularly informative set of variants to study processes of adaptation in human evolution. Future studies with larger datasets and improved methods will further facilitate this line of research and improve our understanding of adaptive processes in human history.

## Methods

### Gene-environment interaction and selection datasets

We leveraged information on GxE loci and contributing environmental factors from Namba et al.^17^. We considered all GxE variants identified in the UK Biobank screen of that study that were either directly tested or for which we could identify a high-LD proxy (r^2^> 0.95) in the Ancient Genome Selection (AGES) database^18^. To identify proxy variants, we downloaded LD matrices for the UK Biobank cohort from https://registry.opendata.aws/ukbb-ld (accessed March-May 2026). In general, LD values between variant pairs were derived from these UK Biobank LD matrices. The only exception was the analysis of LD between GxE variants and local selection lead SNPs, for which we used LD information from the AGES cohort, which was based on European samples from the 1,000 Genomes Project^45^. Additionally, GxE variants located within the Major Histocompatibility Complex were excluded from the analysis, due to the complex LD structure in this region. Information from both datasets was available in human reference genome build hg19.

Within the GxE dataset by Namba et al, ten of the loci we defined were composed of two variants, associated with a total of nine traits. We quantified the overlap of environments within each locus using the Jaccard index, defined as the number of shared items divided by the total number of unique items across two sets. The average index value across the ten loci was 61.5%. To assess whether this overlap exceeded random expectation, we generated 100,000 sets of ten variant pairs drawn from the original set of 55 GxE variants analyzed in this study and calculated the average Jaccard index for each set^46^. Two-sided P values were then computed based on the permutation results.

### Trait and environmental variable enrichment analyses

We tested for both trait and environment enrichment among the four GxE candidates with signatures of selection using Fisher’s exact test. We performed these analyses at both the variant and locus level. Variant-level tests included all GxE variants considered in our analysis, whereas locus-based tests grouped variants into loci defined by LD (r^2^> 0.5). For trait enrichment at the locus level, a locus was considered associated with a given environment if at least one variant within the locus showed such an association. For the environmental factor “meat and cheese”, we additionally tested the probability of finding the observed pattern of all effect sizes among our candidates having the same direction. This was assessed using a permutation test with 100,000 permutations, with two-sided P values calculated based on observations matching or exceeding the extremity observed in the empirical data. The sign of the interaction effect size was polarized to the derived allele effect.

### GWAS overlap of GxE variants

We downloaded GWAS information for 35 (of 47) traits analyzed by Namba et al^17^ for which summary statistics were available in the Neale Lab UK Biobank Round 2 release (http://www.nealelab.is/uk-biobank/). Most missing traits were disease phenotypes in the GxE study that were defined using combined diagnosis variables, which were generally not tested in the same form by the Neale Lab. As a result, matching GWAS data could not be obtained, and these traits were excluded from our analysis. We assessed the level of polygenicity by collapsing genome-wide significant variants (P<5×10^−8^) that are in LD into individual loci. This procedure was performed iteratively, starting by sorting all variants by P value and selecting the most significant variant as the lead. We then identified all variants with r^2^>0.1 to this lead variant and expanded the set by including any additional variants in LD (r^2^>0.1) with the growing set in successive steps. This process was repeated until no further variants could be added, at which point the set was defined as a locus. The procedure was then repeated on the remaining variants until no genome-wide significant variants remained. We used the number of loci as our polygenicity estimate. Next, for each locus, we tested for signatures of selection. A locus was considered to show evidence of selection if at least one variant in LD with the lead variant (r^2^>0.5) had a selection FDR<0.05 in the AGES database. Finally, for each of the three traits associated with a GxE variant with signatures of selection, we tested for enrichment of GxE variants with selection signatures among genome-wide significant GWAS loci that also showed evidence of selection. For this analysis, we only considered GxE variants with significant GWAS P values. Enrichment was tested within each trait using Fisher’s exact test, and across all three traits using a Mantel-Haenszel test.

### Allele frequency reconstruction using CLUES

Additionally, we reconstructed the allele-frequency trajectories of our GxE candidates across various present-day human populations, leveraging an ancestral recombination graph-based approach previously implemented by Pankratov et al^47^. First, we used Relate (version 1.2.4)^48^ to construct local trees across the genomes of 2,271 unrelated individuals from the 1,000 Genomes Project^49^. We randomly selected 90 samples per population, unless fewer samples were available, in which case we included all samples (ASW: 61, MXL: 64, MSL & PEL: 85, BEB: 86). We used Relate with default parameters and auxiliary files for human reference hg38, provided with the tool: a pilot mask to remove positions prone to inaccessibility for short-read sequencing technologies, a recombination map, and the human ancestral genome. Then, we extracted subtrees and estimated coalescent rates for each of the 26 1,000 Genomes populations. Lastly, for three of the four GxE candidate variants that passed the Relate quality filters, we extracted its corresponding tree and generated 200 trees with re-estimated branch lengths. These trees and coalescent rates were used as input for CLUES^50^, which infers selection statistics and allele frequency trajectories. We ran CLUES for each GxE candidate variant across all populations over the last 500 generations using a published computational pipeline^51^. Since there is an uncertainty in branch length computation, we repeated branch resampling and selection estimation analysis 5 times per tested variant. Due to differences in human reference assemblies between the GxE and AGES datasets (hg19) and the high-coverage 1,000 Genomes project data (hg38), we converted genomic coordinates from hg19 to hg38 using LiftOver^52^. In total, we analyzed 69 variant-population pairs (Urate-locus1: 22 pairs, Urate-locus2: 21 pairs, TC-locus3: 26 pairs), excluding pairs for which a given variant was not detected in the population (Supplementary table 8).

Frequency trajectories were defined by assigning, at each time point, the frequency with the highest posterior probability. Next, we excluded allele frequency trajectories that showed high levels of divergence with other trajectory replicates within a given variant-population pair. Specifically, we calculated the mean absolute difference to the other four trajectories across all time points (500 generations) and then averaged these values to obtain a per-trajectory distance measure. To identify outlier trajectories, we calculated a distance-based measure for each trajectory across all variant-population pairs and defined a threshold using the overall distribution of these values. Trajectories were classified as outliers if their distance exceeded the median by more than three median absolute deviations. In total, we identified 26 outlier trajectories (7.5%). In three instances, more than two outlier trajectories were associated with the same variant-population pair. Due to the high levels of divergence across trajectories within these pairs, we considered the reconstructed trajectories unreliable and excluded the three pairs from further analysis. Next, we generated one trajectory for each variant-population pair, by averaging the allele frequencies for trajectory replicates across time points (Supplementary table 8).

### Allele frequency breakpoint estimator

To assess whether the allele frequency trajectories of our four GxE candidates showed differences over time, including periods of increased or decreased rates of change, we performed a test to identify potential trajectory breakpoints. Breakpoint detection was performed using a piecewise linear modeling framework. For each trajectory, a single linear regression fitted across all time points was compared to a two-segment model composed of two independent linear regressions separated by a candidate breakpoint. For all possible breakpoint locations, the trajectory was partitioned into two segments and each segment was fitted independently. Model fit was evaluated using the residual sum of squares (RSS). The optimal breakpoint was defined as the time point that maximized the improvement in fit of the two-line model relative to the single-line model. This improvement (“gain”) was calculated as the difference between the RSS of the single-line model and the combined RSS of the two segments. To enable comparison across trajectories, the gain was normalized by the RSS of the single-line model, resulting in a relative gain estimate that reflects the proportion of variance explained by introducing a breakpoint. Statistical significance of the model improvement was assessed using a permutation-based approach. For each trajectory, a null distribution of maximal gain values was generated by randomly permuting the frequency values while preserving the time structure, thereby removing temporal dependence. For each permuted dataset, the breakpoint detection procedure was repeated and the maximal gain recorded. The P value was estimated as the proportion of permuted gains that were greater than or equal to the observed gain, providing a non-parametric measure of significance (Supplementary table 8). We applied this approach to both the CLUES and empirical AGES frequency trajectories.

### Molecular and phenotypic annotation of GxE candidate variants

We used the Ensembl Variant Effect Predictor (VEP) web interface for the human genome build hg19 (GRCh37) 53 to annotate the putative molecular effects of the four candidate GxE variants (Supplementary table 7). In addition, we included all variants in high LD (r^2^>0.5 in the UK Biobank) with these candidates. Furthermore, we examined trait associations for these variants in the Open Targets database 54 (version March 2026 26.03) and included all GWAS hits with genome-wide significance (P<5×10^−8^) that were assigned to a credible set, i.e. a set of SNPs associated with a local association peak.

## Acknowledgements

This work was supported by the Ministry of Education and Research Centres of Excellence grant TK214 “Centre of Excellence for Personalised Medicine” and the Estonian Research Council grant PRG3113. L.W. was supported through the Erasmus+ programme (project code: 2025-1-DE01-KA131-HED-000333845). Data analyses were carried out in part in the High-Performance Computing Center of the University of Tartu. Finally, we thank the members of the Center of Genomics, Evolution and Medicine in Tartu and the Archaeo- and Paleogenetics Group in Tübingen for valuable discussions and feedback on this project. This preprint was typeset with the bioRxiv word template by @Chrelli: www.github.com/chrelli/bioRxiv-word-template

## Author contributions

Lotte Wijnhoven (Data curation [lead], Formal analysis [lead], Investigation [lead], Methodology [supporting], Visualization [lead], Writing-original draft [supporting], Writing-review & editing [supporting]), Danat Yermakovich (Data curation [supporting], Formal analysis [supporting], Writing-review & editing [supporting]), Michael Dannemann (Conceptualization [lead], Data curation [supporting], Formal analysis [supporting], Investigation [lead], Methodology [lead], Project administration [lead], Supervision [lead], Visualization [lead], Writing-original draft [lead], Writing-review & editing [lead]).

## Data availability

All data needed to evaluate the conclusions in the paper are present in the paper and/or the Supplementary Materials.

## Code availability

The code used to perform the analyses described in this study is publicly available on GitHub under github.com/lottewijnhoven/master_thesis_project.

## Supplementary Material

### Supplementary figure legends

Supplementary figure 1: Empirical and modelled allele frequency trajectories of four GxE candidate variants.

**(A-D)** The empirical (blue) and modelled (orange) derived allele frequency trajectories over the last 10,000 years in the AGES cohort are shown as solid lines. Confidence intervals are indicated by shaded areas in the corresponding colors. Dashed blue lines indicate change points that partition each trajectory into two segments with the greatest difference in allele frequency change. The relative gain and P value associated with the change-point test are shown below the x-axis.

Supplementary figure 2: Trait and environment associations for GxE variants.

**(A-J)** For each environment **(A-H)** and trait **(I-J)** associated with at least one of the four candidate GxE variants, but not enriched for them, GxE variants are shown as squares and partitioned into two sets: one comprising all four candidate variants (upper set) and a second comprising all 51 variants without signatures of selection (lower set). Variants within both sets are ordered according to their genomic position, and variant pairs belonging to the same locus are connected by a black line. Variants associated with the focal environment or trait are highlighted in blue (environments) or salmon (traits), respectively.

Supplementary figure 3: Reconstructed derived allele frequency trajectories for three of the four candidate GxE variants using CLUES.

For each combination of one of the three tested GxE variants (shown as the rsID at the top of each panel) and a 1,000 Genomes population in which the variant was tested, the average reconstructed derived allele frequency trajectory across replicates over the last 500 generations is shown as a solid black line. In addition, the estimated change point for each variant-population combination is indicated by a vertical black line. The relative gain and P value associated with the change point test are shown below the x-axis.

Supplementary figure 4: Polygenicity and selection information for GxE study traits.

**(A)** For each trait, the number of independent significant GWAS loci (x-axis) and the number of independent significant GWAS loci containing at least one variant with a selection FDR<0.05 that is in LD (r^2^>0.5) with the corresponding GWAS lead SNP (y-axis) are shown. Traits associated with candidate GxE variants are shown in green, whereas traits not associated with candidate GxE variants are shown in blue. Circle size is proportional to the number of GxE variants associated with each trait. **(B)** The proportion of independent loci with selection signatures among all independent loci is shown for each trait, ordered by this proportion. Traits associated with candidate GxE variants are shown in light blue, whereas all other traits are shown in blue. Information on the trait abbreviations used on the y-axis can be found in supplementary table 1.

### Supplementary table legends

Supplementary table S1: General information as well as polygenicity and selection estimates for traits analyzed by Namba et al in the UK Biobank.

Supplementary table S2: Information about the environmental interaction factors included in the analysis of the UK Biobank cohort by Namba et al.

Supplementary table S3: Selection and GWAS information for GxE variants identified by Namba et al in the UK Biobank. Supplementary table S4: Trait and environment enrichment results for GxE with signatures of selection.

Supplementary table S5: Derived allele frequency distribution of candidate GxE variants in all 26 populations from the 1,000 Genomes cohort.

Supplementary table S6: Overlap of GxE candidate variants with credible GWAS sets from the OpenTargets database. Supplementary table S7: Predicted molecular effects of candidate GxE variants and of variants in high LD with them.

Supplementary table S8: Reconstructed allele frequency trajectories and change point statistics for three candidate GxE variants across 1,000 Genomes populations.

