## Supplementary figures and images for "Gene-environment interaction loci with signatures of mid- to late Holocene selection show evidence for diet-dependent genetic effects"

### Supplementary figure S1

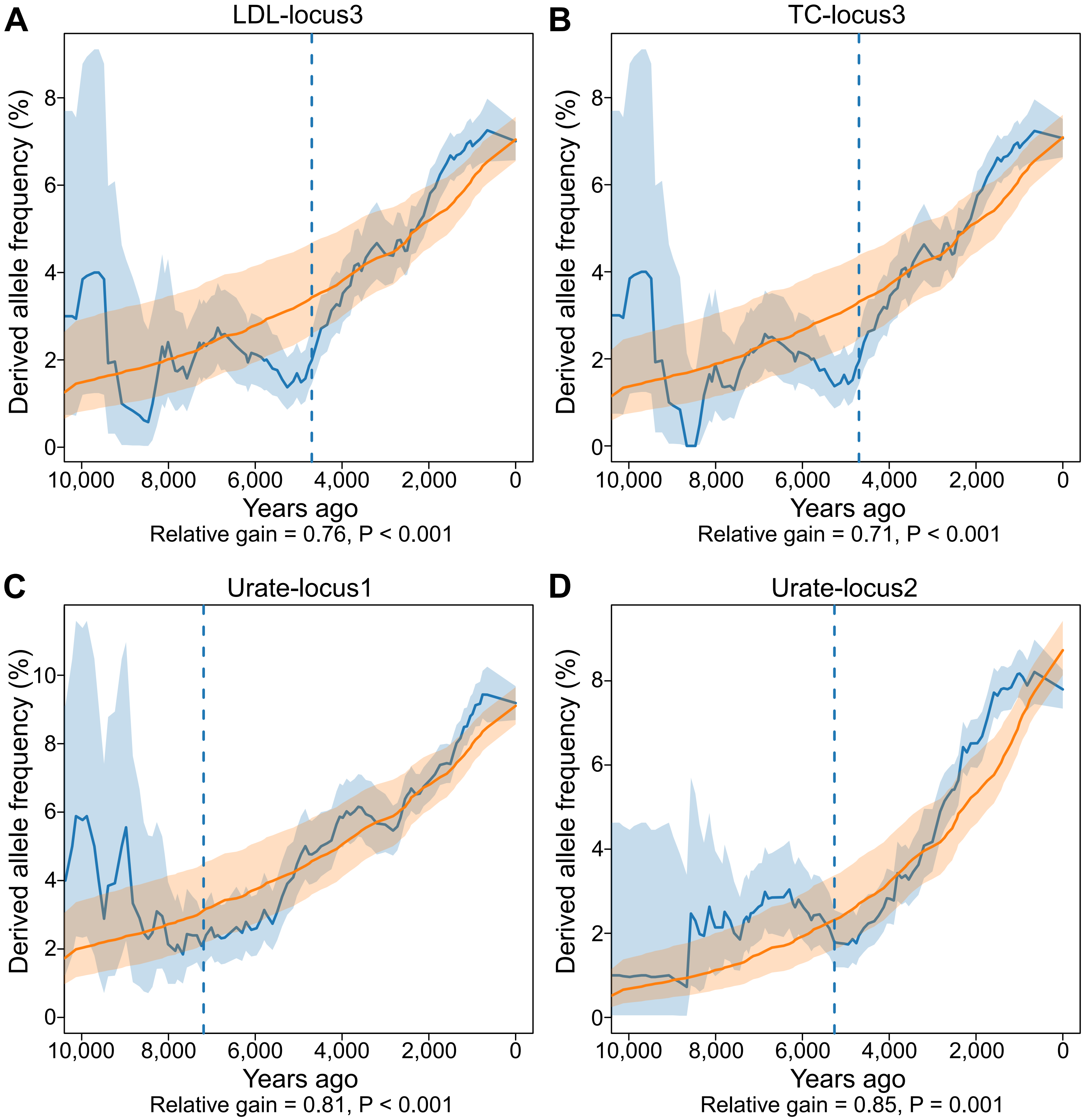

### Supplementary figure S2

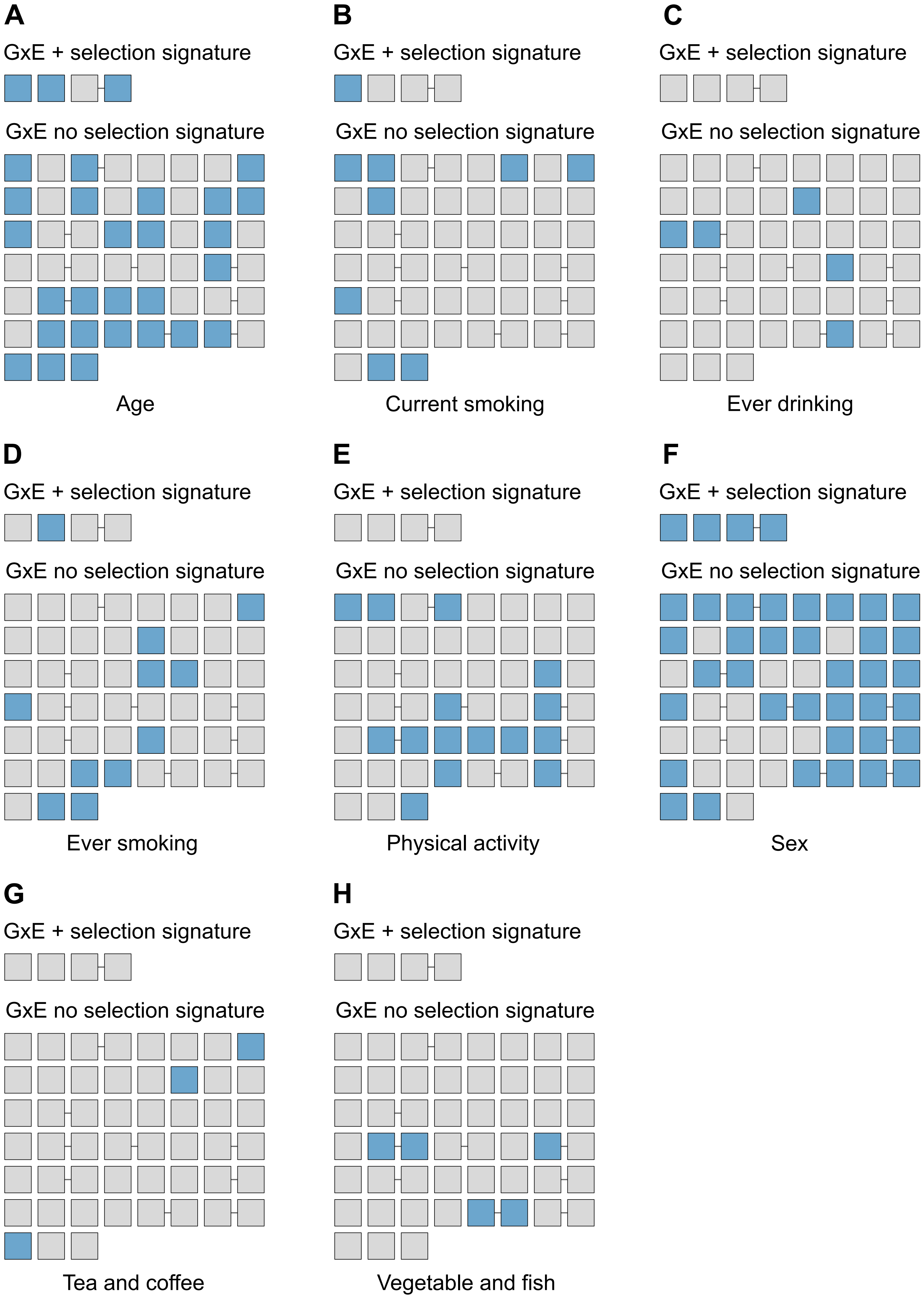

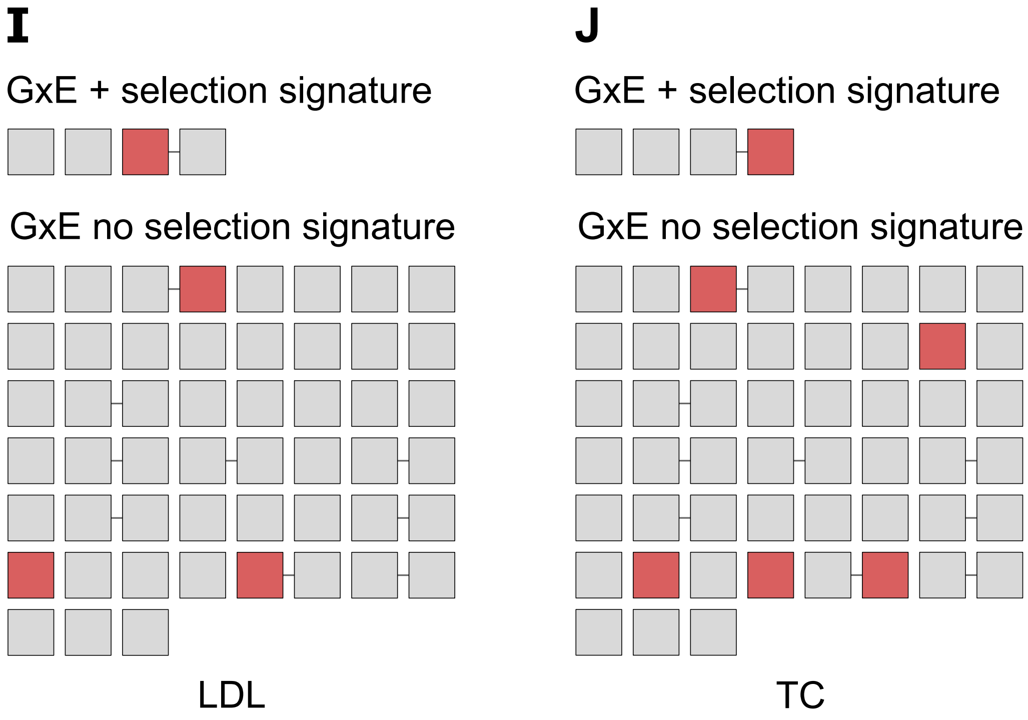

### Supplementary figure S4

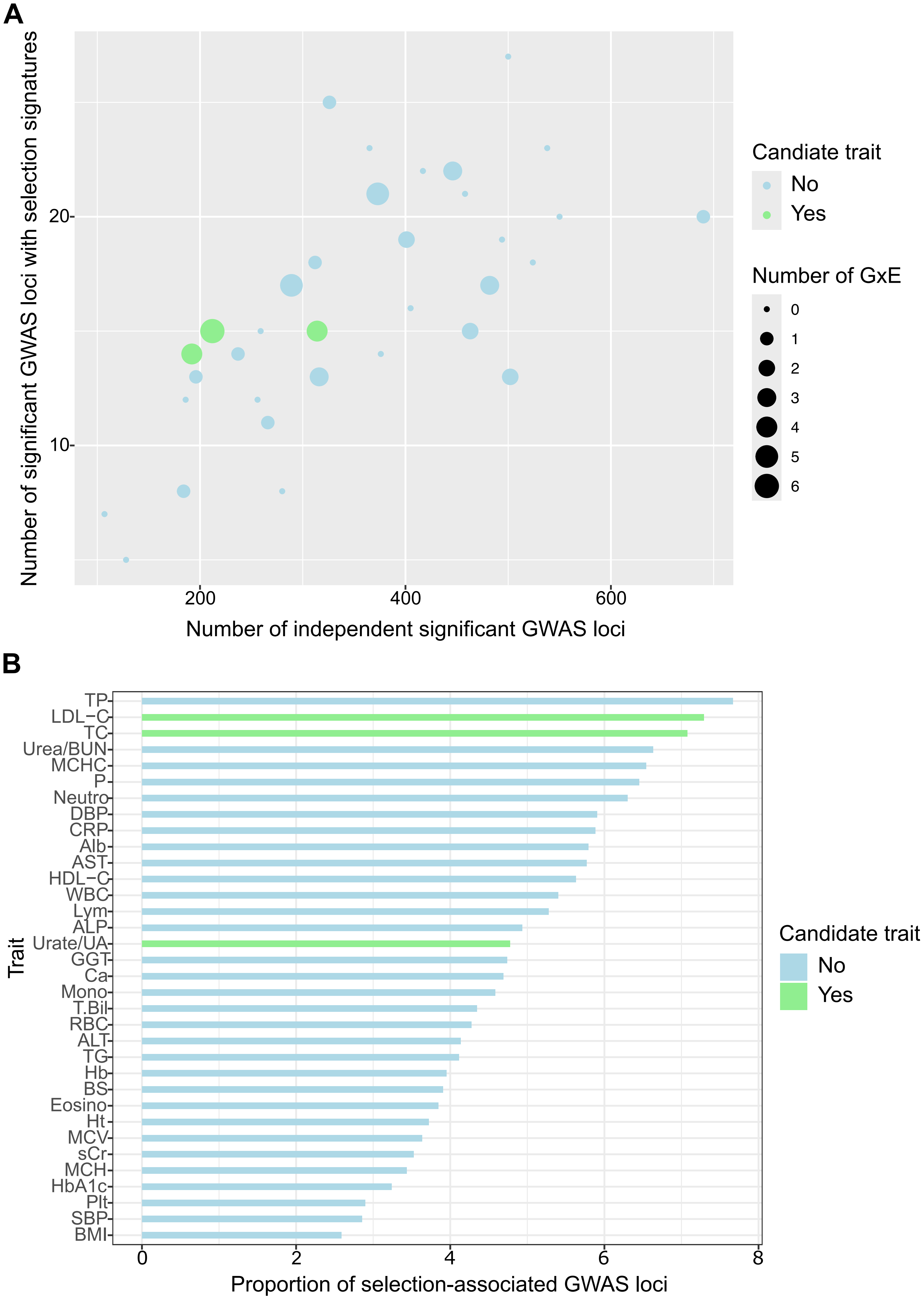
