## Supplementary figure S3 for "Gene-environment interaction loci with signatures of mid- to late Holocene selection show evidence for diet-dependent genetic effects"

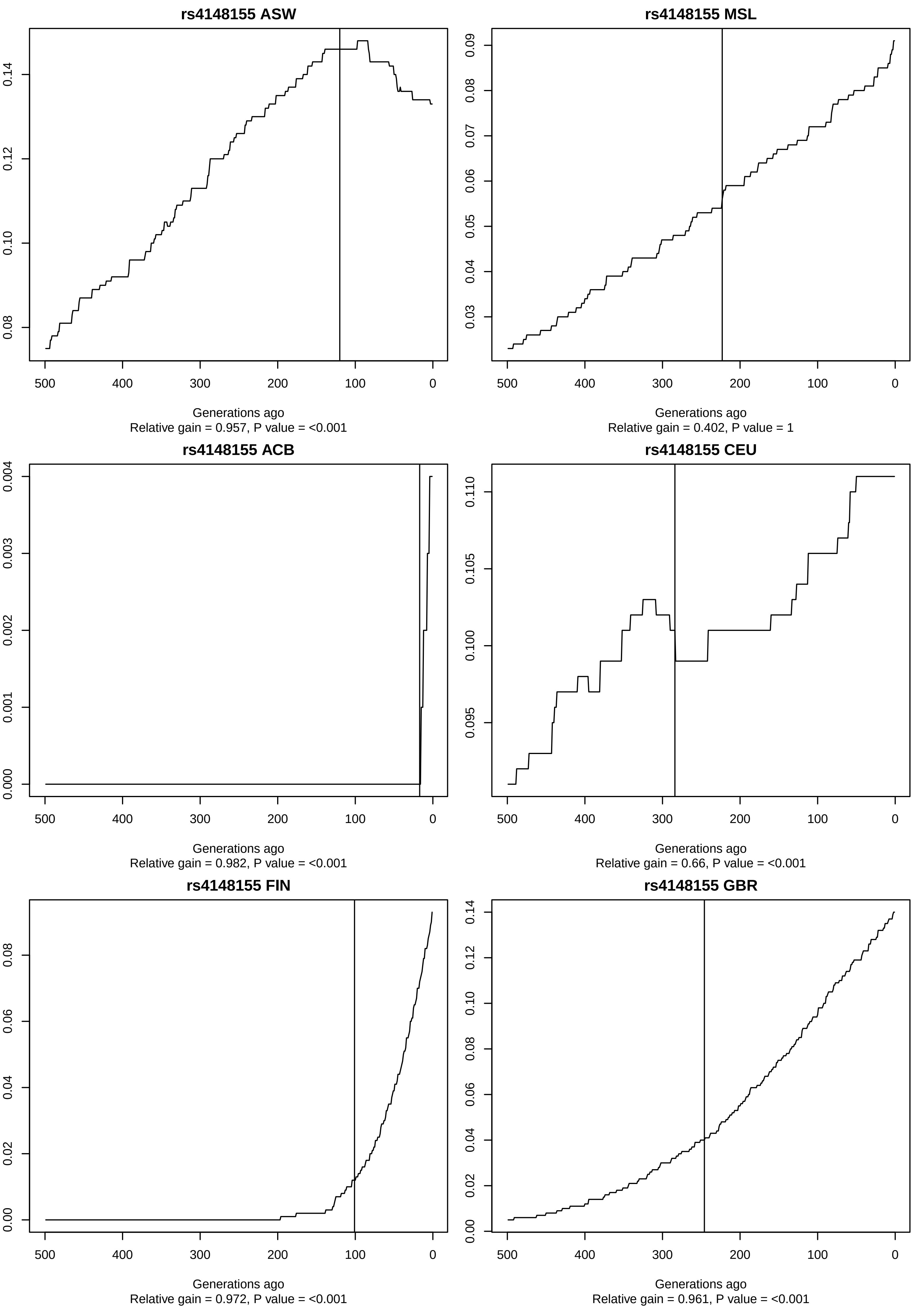
rs4148155 ASW
Generations ago
Relative gain = 0.957, P value = <0.001
rs4148155 MSL
Generations ago
Relative gain = 0.402, P value = 1
rs4148155 ACB
Generations ago
Relative gain = 0.982, P value = <0.001
rs4148155 CEU
Generations ago
Relative gain = 0.66, P value = <0.001
rs4148155 FIN
Generations ago
Relative gain = 0.972, P value = <0.001
rs4148155 GBR
Generations ago
Relative gain = 0.961, P value = <0.001

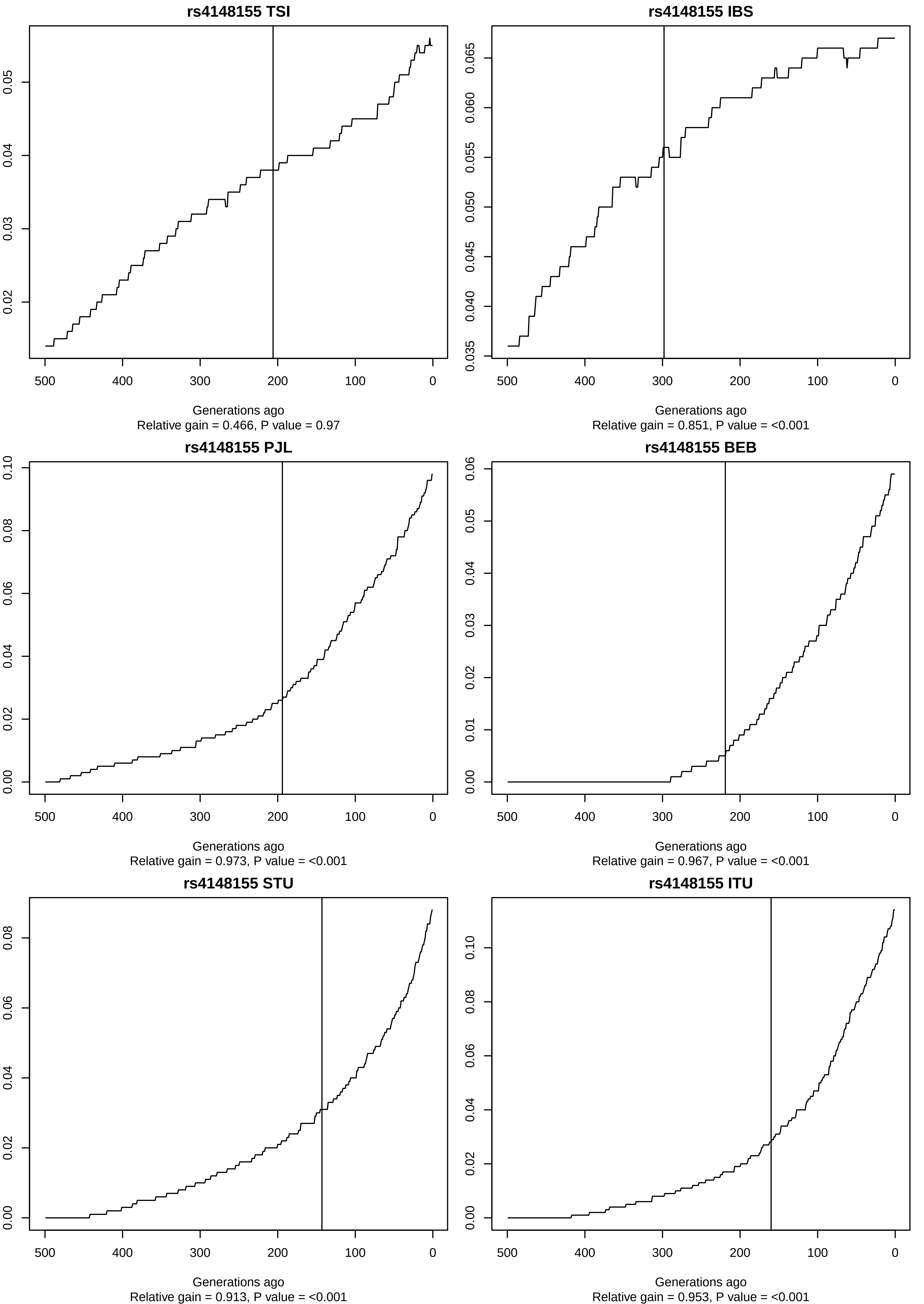
rs4148155 TSI
Generations ago
Relative gain = 0.466, P value = 0.97
rs4148155 IBS
Generations ago
Relative gain = 0.851, P value = <0.001
rs4148155 PJL
Generations ago
Relative gain = 0.973, P value = <0.001
rs4148155 BEB
Generations ago
Relative gain = 0.967, P value = <0.001
rs4148155 STU
Generations ago
Relative gain = 0.913, P value = <0.001
rs4148155 ITU
Generations ago
Relative gain = 0.953, P value = <0.001

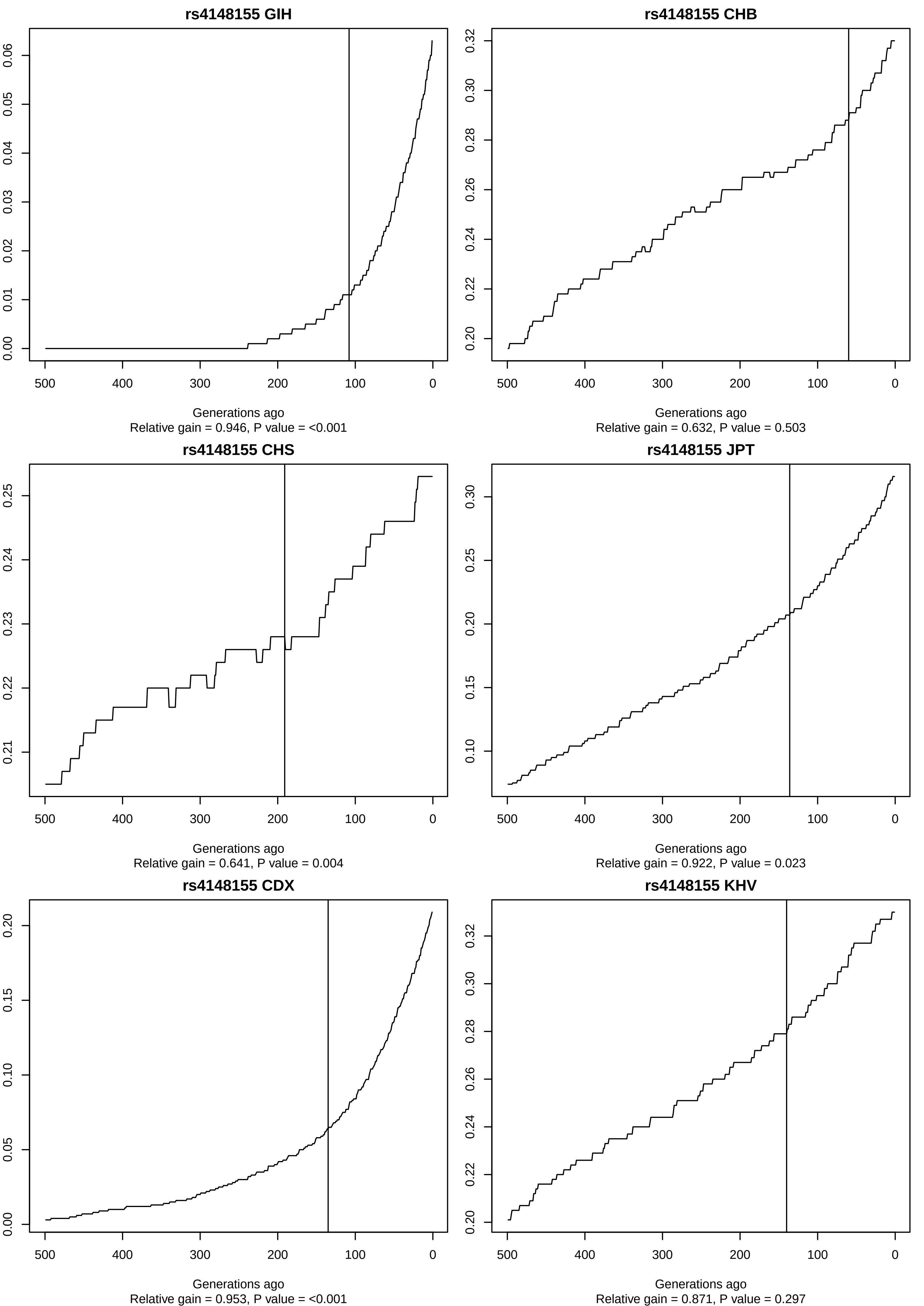

rs4148155 GIH
Generations ago
Relative gain = 0.946, P value = <0.001
rs4148155 CHB
Generations ago
Relative gain = 0.632, P value = 0.503
rs4148155 CHS
Generations ago
Relative gain = 0.641, P value = 0.004
rs4148155 JPT
Generations ago
Relative gain = 0.922, P value = 0.023
rs4148155 CDX
Generations ago
Relative gain = 0.953, P value = <0.001
rs4148155 KHV
Generations ago
Relative gain = 0.871, P value = 0.297

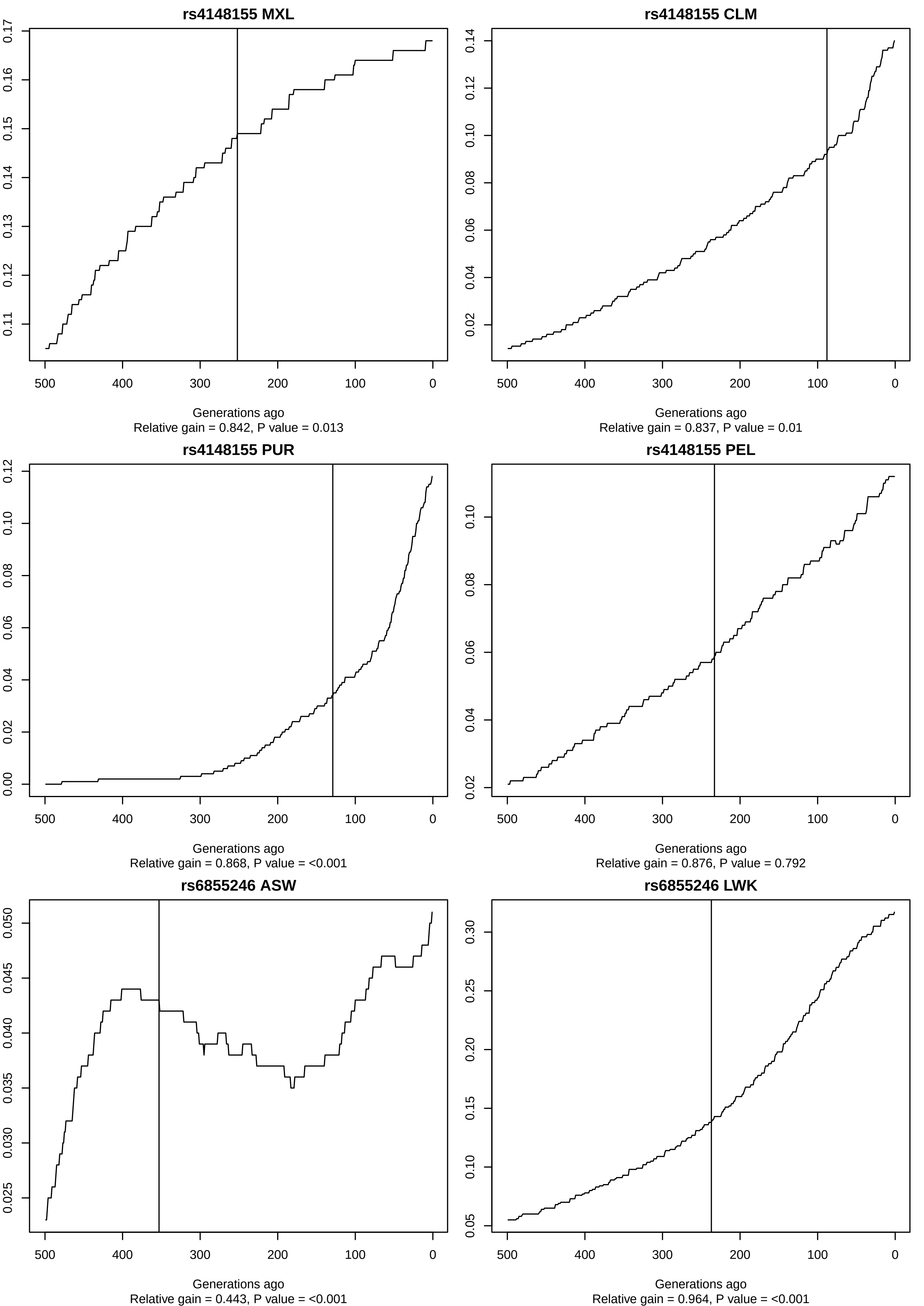
rs4148155 MXL
Generations ago
Relative gain = 0.842, P value = 0.013
rs4148155 CLM
Generations ago
Relative gain = 0.837, P value = 0.01
rs4148155 PUR
Generations ago
Relative gain = 0.868, P value = <0.001
rs4148155 PEL
Generations ago
Relative gain = 0.876, P value = 0.792
rs6855246 ASW
Generations ago
Relative gain = 0.443, P value = <0.001
rs6855246 LWK
Generations ago
Relative gain = 0.964, P value = <0.001

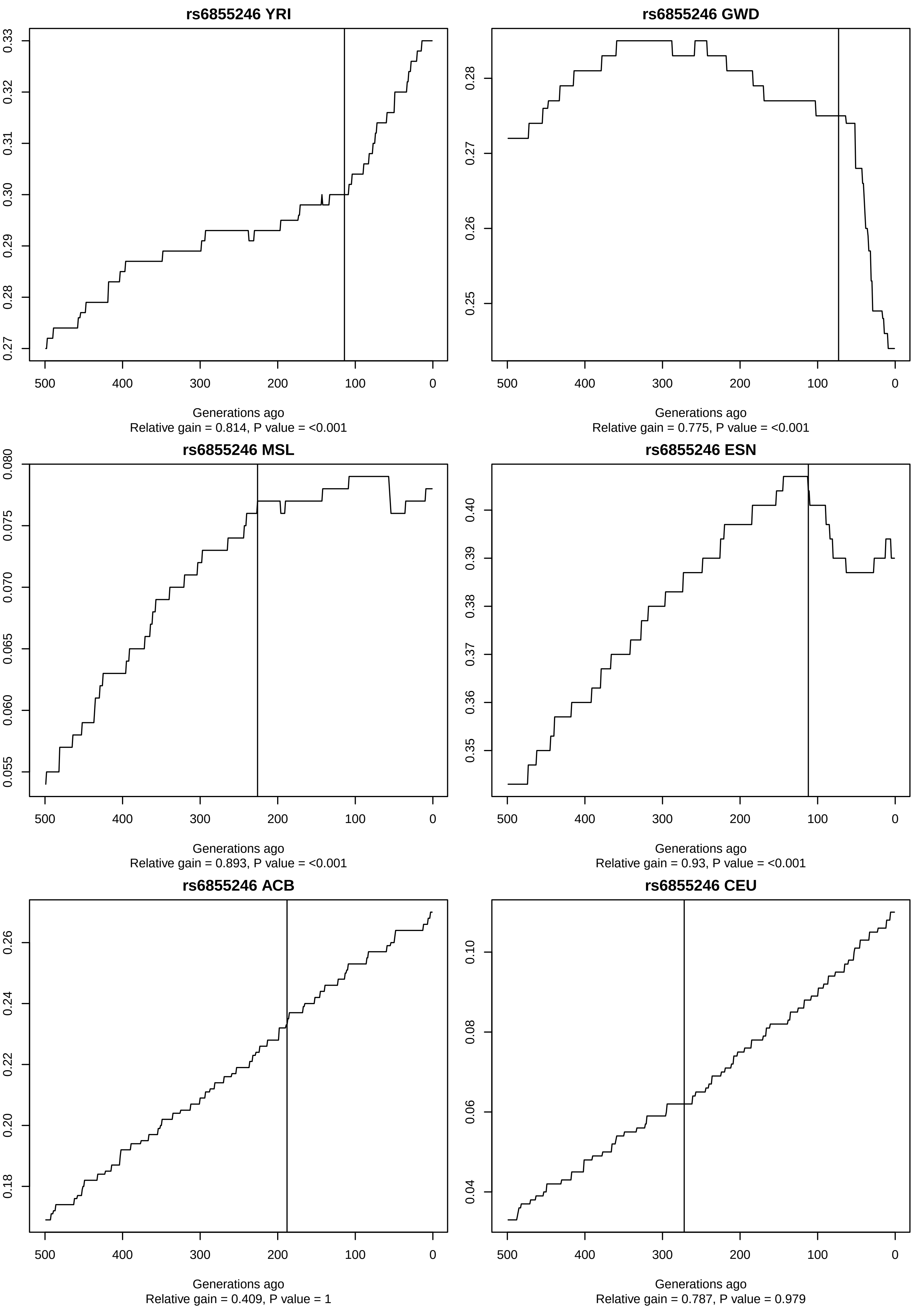
rs6855246 YRI
Generations ago
Relative gain = 0.814, P value = <0.001
rs6855246 GWD
Generations ago
Relative gain = 0.775, P value = <0.001
rs6855246 MSL
Generations ago
Relative gain = 0.893, P value = <0.001
rs6855246 ESN
Generations ago
Relative gain = 0.93, P value = <0.001
rs6855246 ACB
Generations ago
Relative gain = 0.409, P value = 1
rs6855246 CEU
Generations ago
Relative gain = 0.787, P value = 0.979

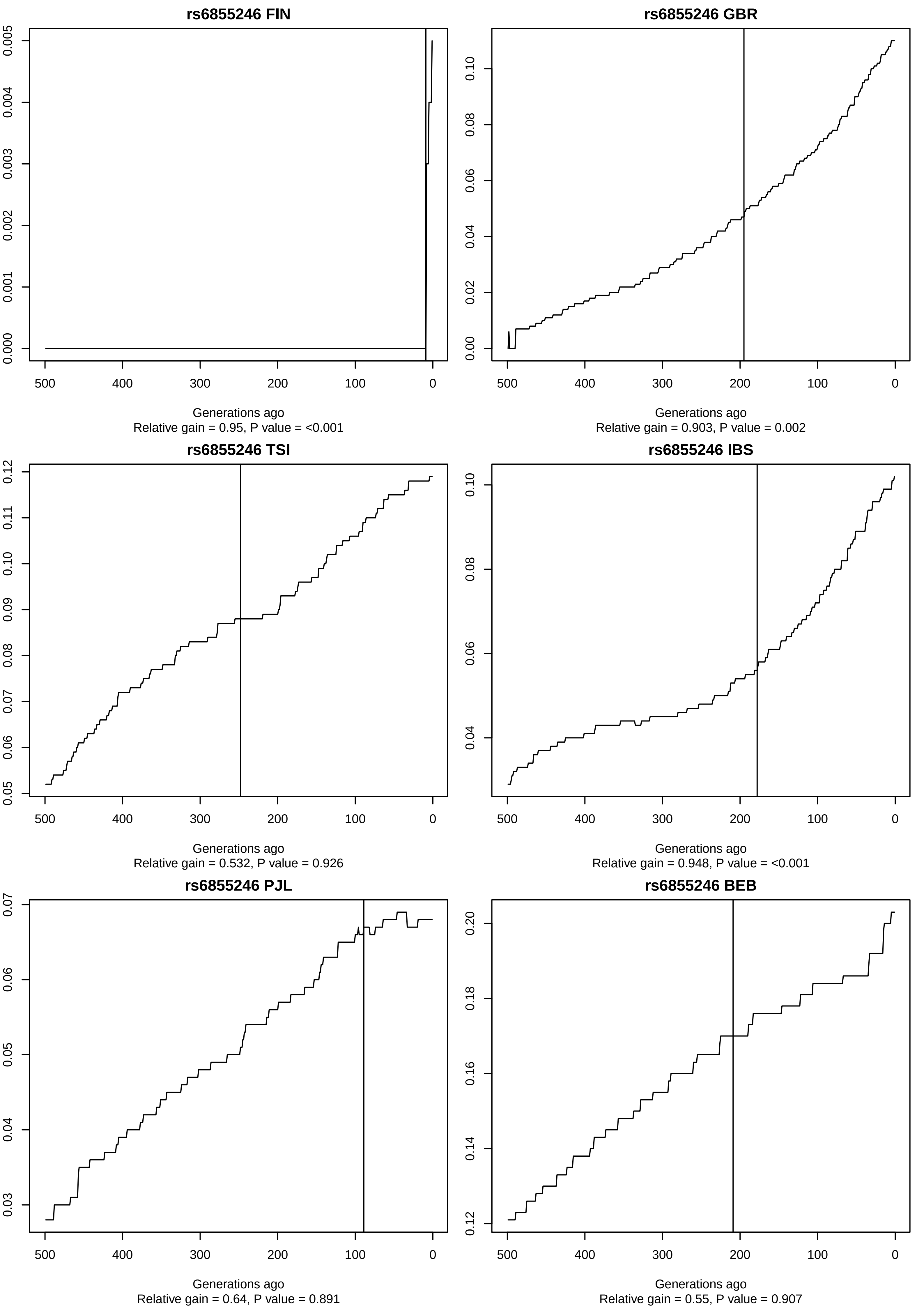
rs6855246 FIN
Generations ago
Relative gain = 0.95, P value = <0.001
rs6855246 GBR
Generations ago
Relative gain = 0.903, P value = 0.002
rs6855246 TSI
Generations ago
Relative gain = 0.532, P value = 0.926
rs6855246 IBS
Generations ago
Relative gain = 0.948, P value = <0.001
rs6855246 PJL
Generations ago
Relative gain = 0.64, P value = 0.891
rs6855246 BEB
Generations ago
Relative gain = 0.55, P value = 0.907

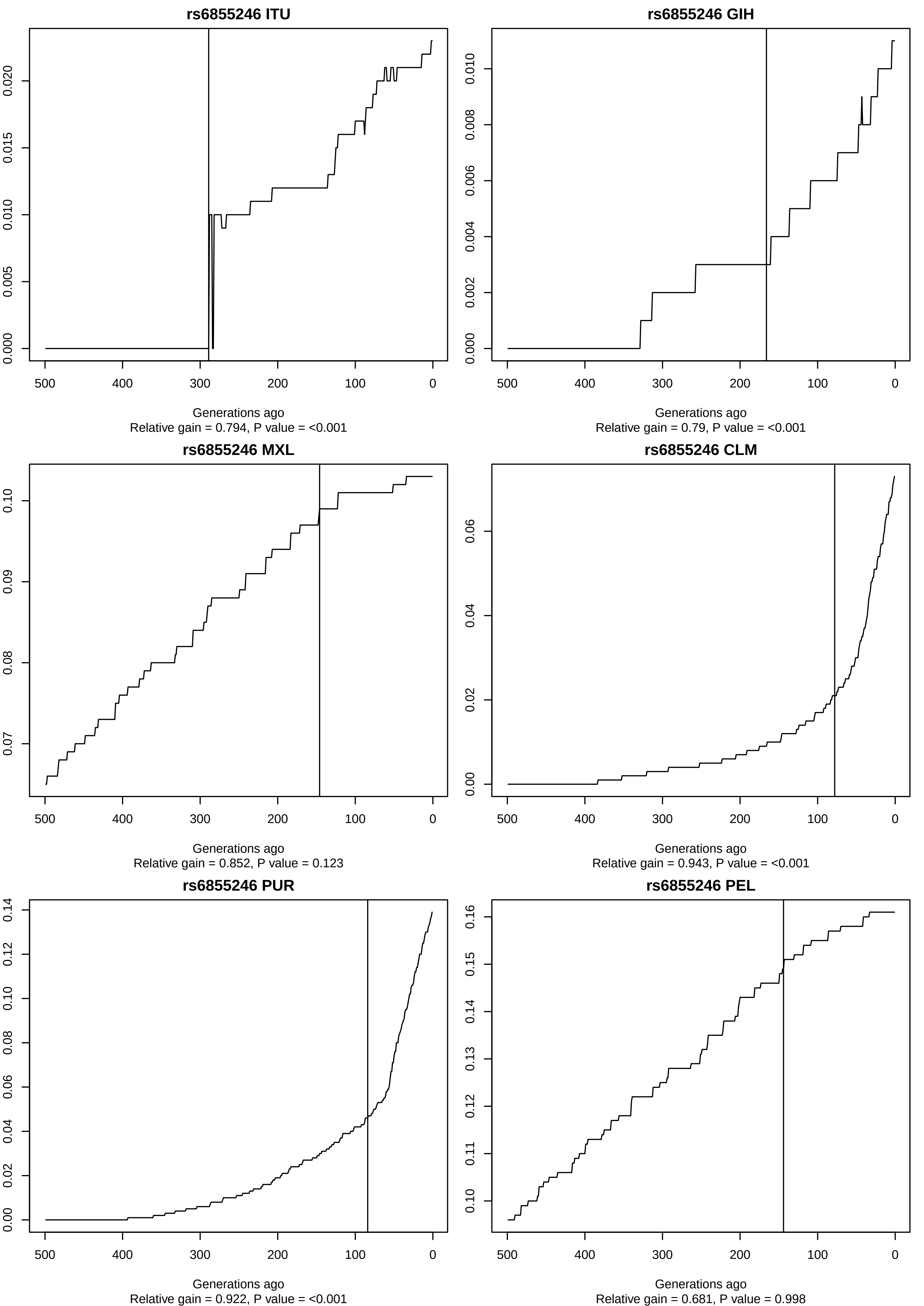
rs6855246 ITU
Generations ago
Relative gain = 0.794, P value = <0.001
rs6855246 GIH
Generations ago
Relative gain = 0.79, P value = <0.001
rs6855246 MXL
Generations ago
Relative gain = 0.852, P value = 0.123
rs6855246 CLM
Generations ago
Relative gain = 0.943, P value = <0.001
rs6855246 PUR
Generations ago
Relative gain = 0.922, P value = <0.001
rs6855246 PEL
Generations ago
Relative gain = 0.681, P value = 0.998

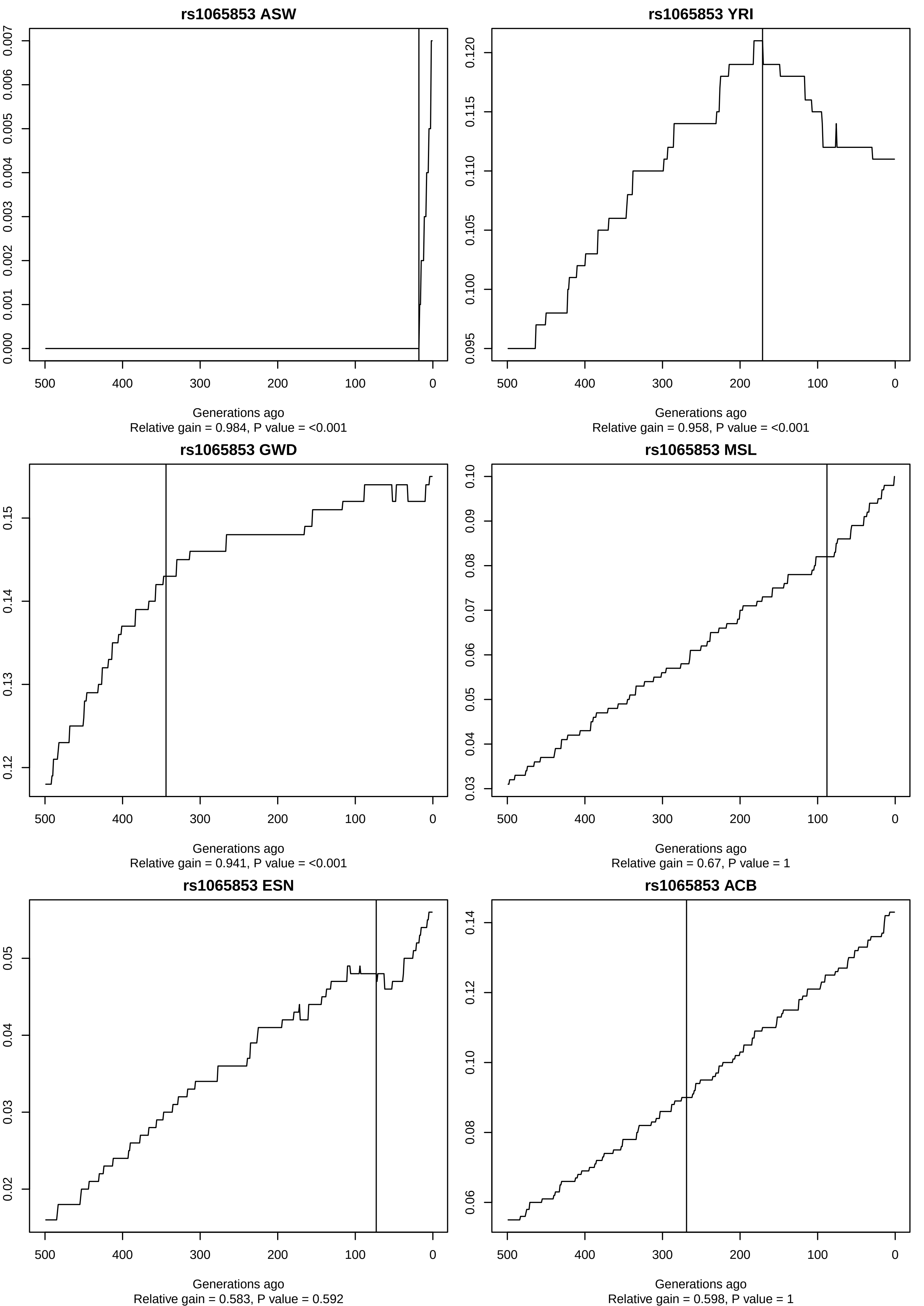
rs1065853 ASW
Generations ago
Relative gain = 0.984, P value = <0.001
rs1065853 YRI
Generations ago
Relative gain = 0.958, P value = <0.001
rs1065853 GWD
Generations ago
Relative gain = 0.941, P value = <0.001
rs1065853 MSL
Generations ago
Relative gain = 0.67, P value = 1
rs1065853 ESN
Generations ago
Relative gain = 0.583, P value = 0.592
rs1065853 ACB
Generations ago
Relative gain = 0.598, P value = 1

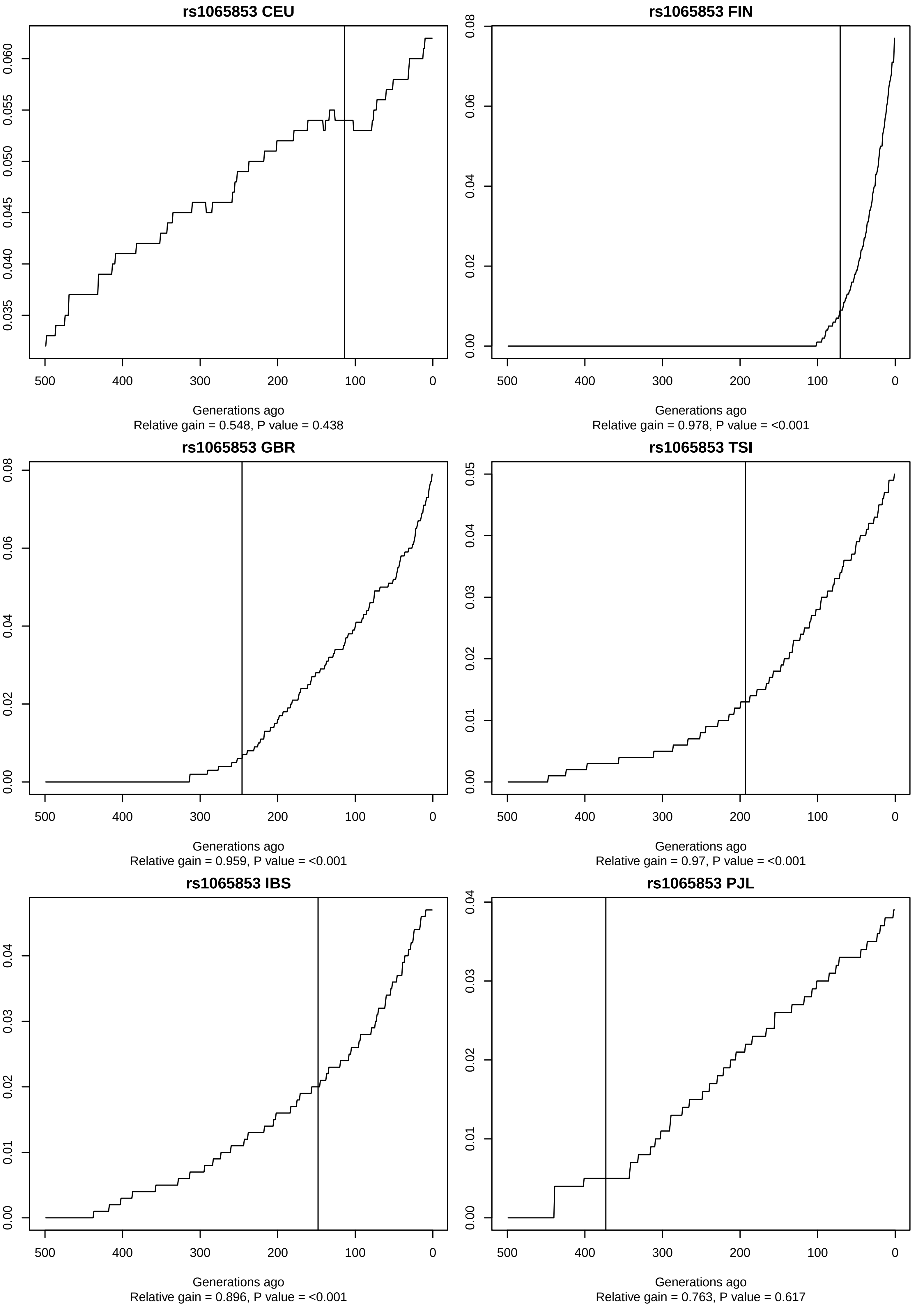
rs1065853 CEU
Generations ago
Relative gain = 0.548, P value = 0.438
rs1065853 FIN
Generations ago
Relative gain = 0.978, P value = <0.001
rs1065853 GBR
Generations ago
Relative gain = 0.959, P value = <0.001
rs1065853 TSI
Generations ago
Relative gain = 0.97, P value = <0.001
rs1065853 IBS
Generations ago
Relative gain = 0.896, P value = <0.001
rs1065853 PJL
Generations ago
Relative gain = 0.763, P value = 0.617

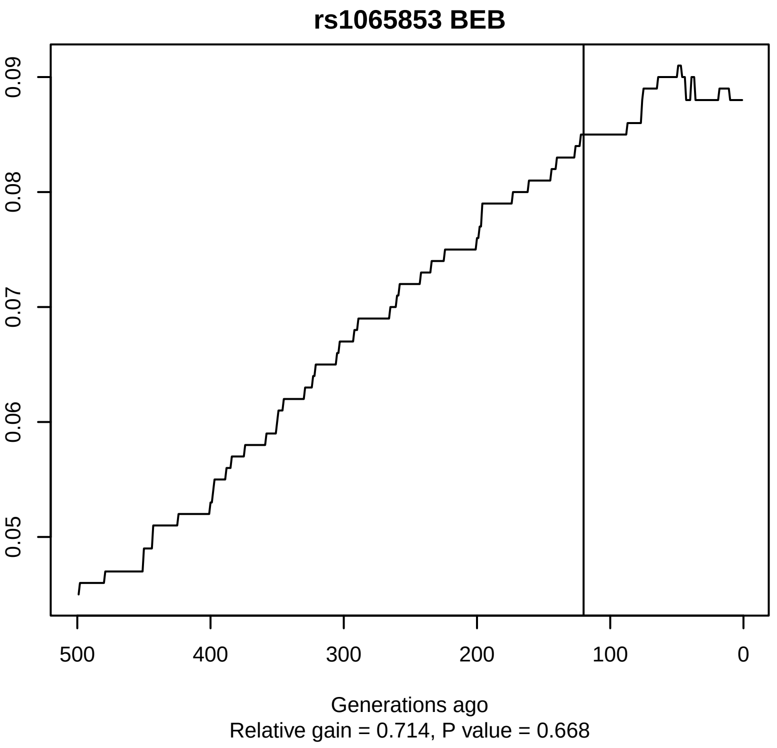
rs1065853 BEB
Generations ago
Relative gain = 0.714, P value = 0.668

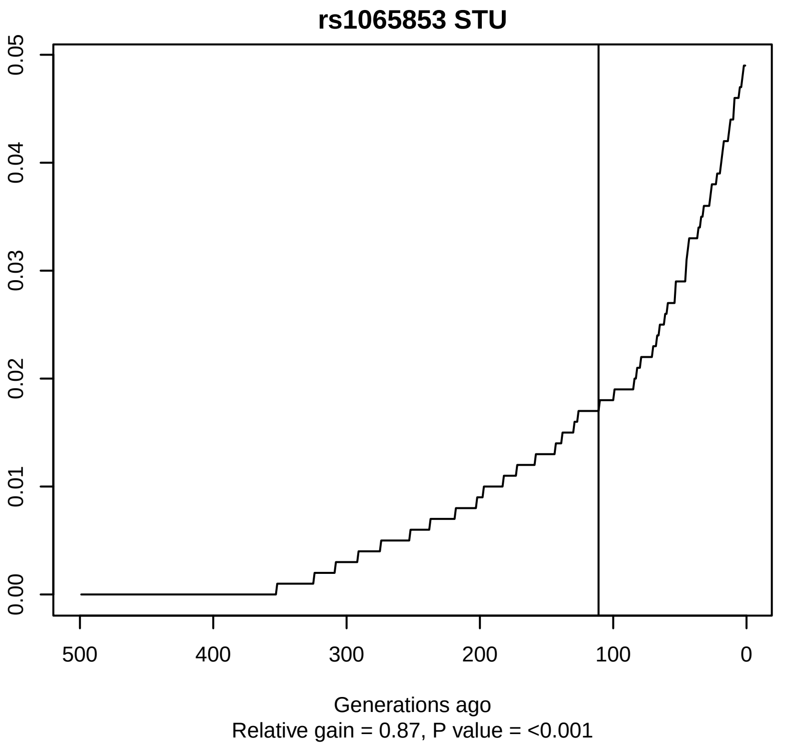
rs1065853 STU
Generations ago
Relative gain = 0.87, P value = <0.001

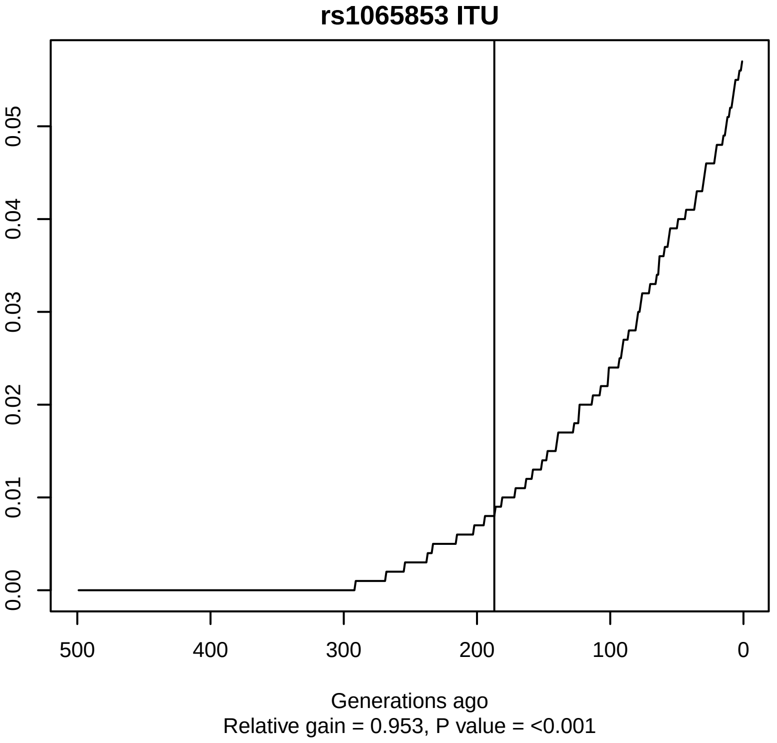
rs1065853 ITU
Generations ago
Relative gain = 0.953, P value = <0.001

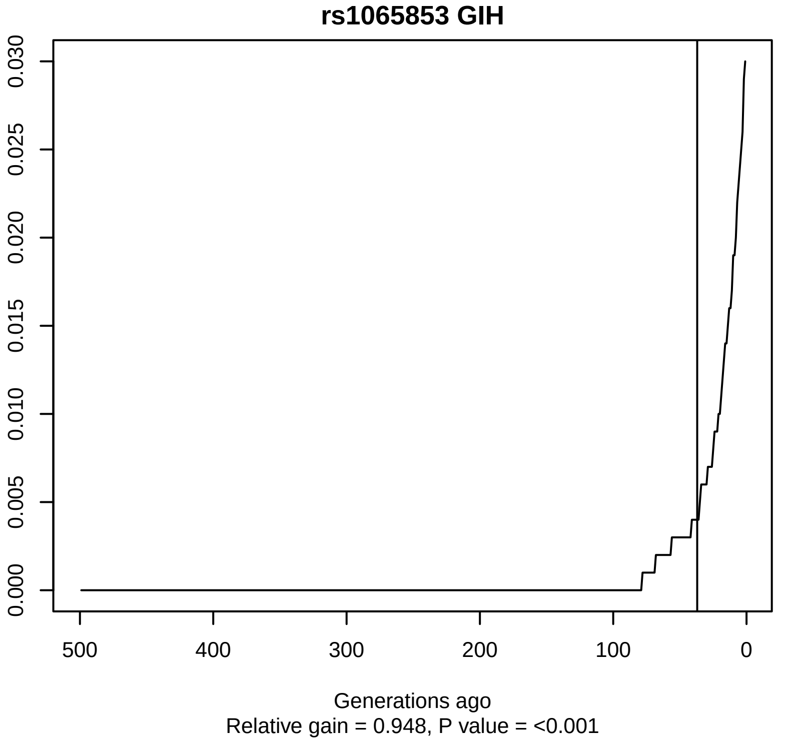
rs1065853 GIH
Generations ago
Relative gain = 0.948, P value = <0.001

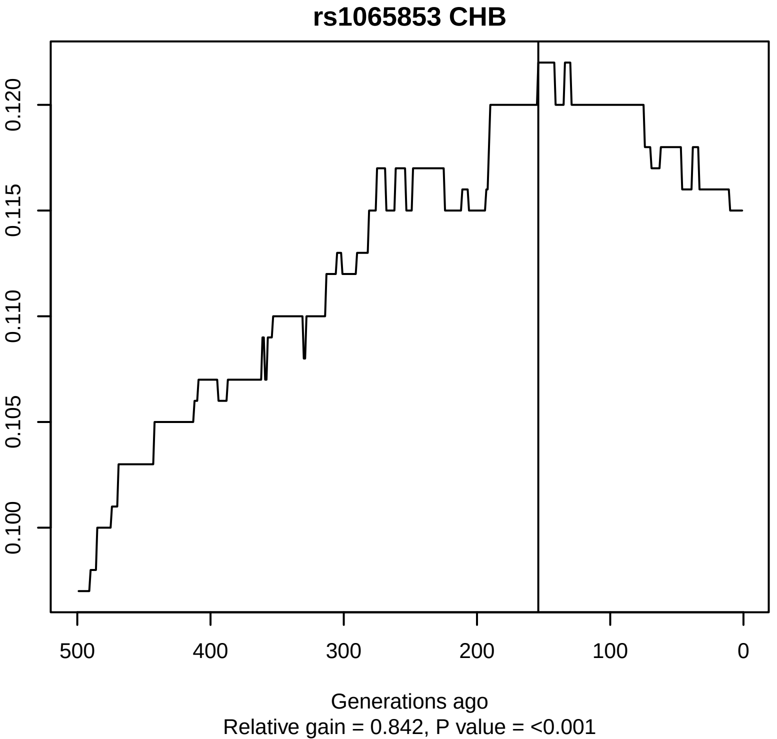
rs1065853 CHB
Generations ago
Relative gain = 0.842, P value = <0.001

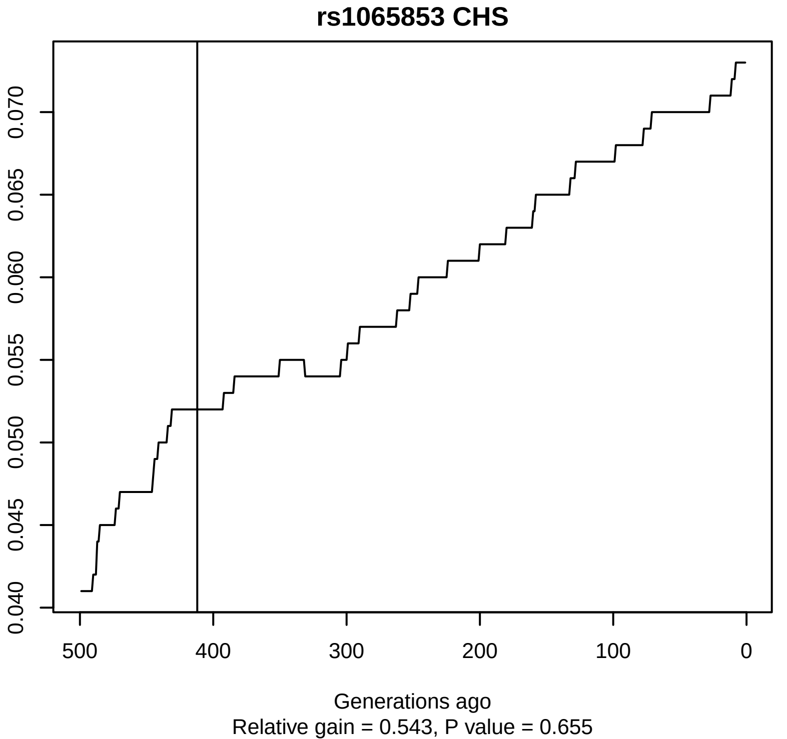
rs1065853 CHS
Generations ago
Relative gain = 0.543, P value = 0.655

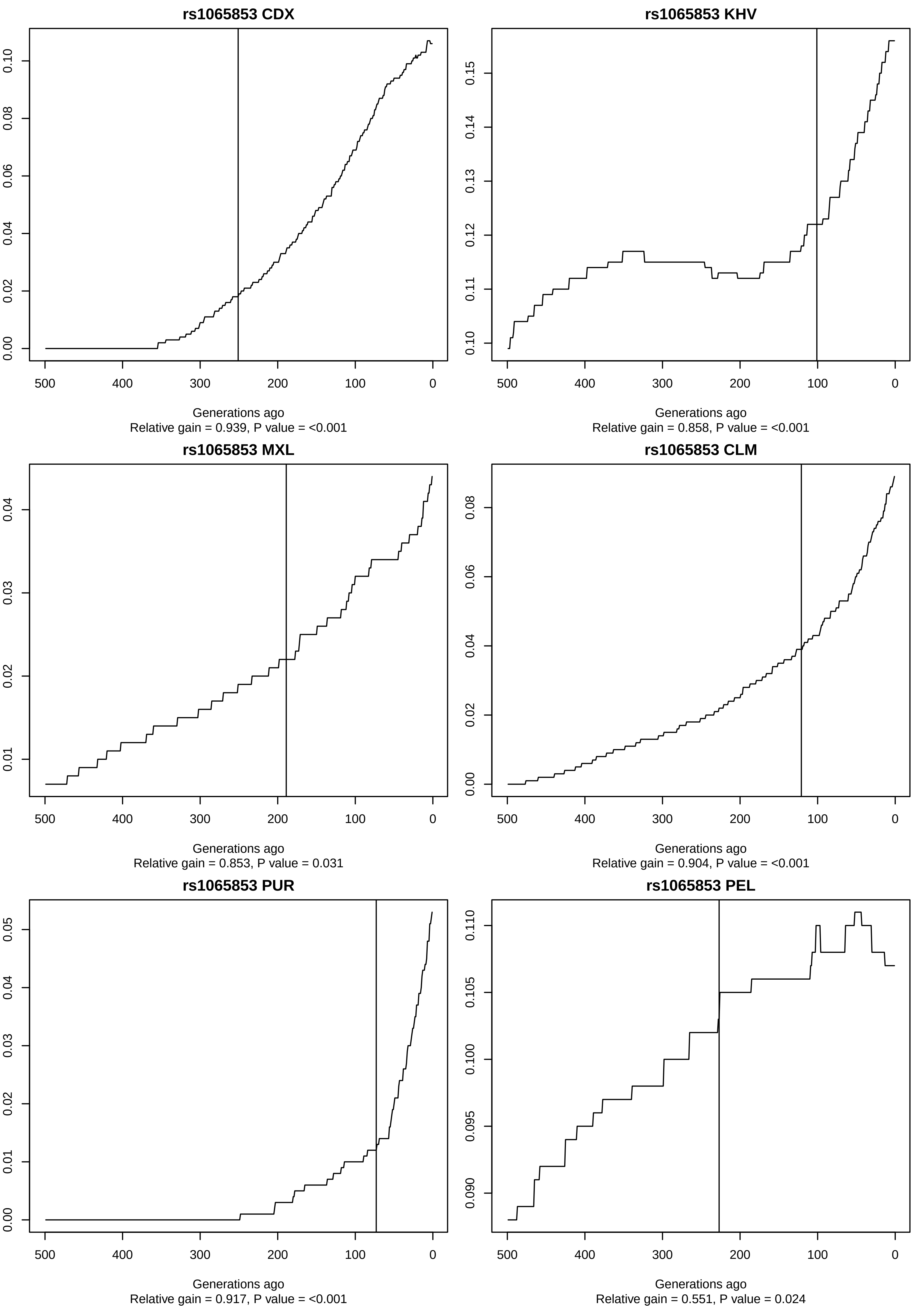
rs1065853 CDX
Generations ago
Relative gain = 0.939, P value = <0.001
rs1065853 KHV
Generations ago
Relative gain = 0.858, P value = <0.001
rs1065853 MXL
Generations ago
Relative gain = 0.853, P value = 0.031
rs1065853 CLM
Generations ago
Relative gain = 0.904, P value = <0.001
rs1065853 PUR
Generations ago
Relative gain = 0.917, P value = <0.001
rs1065853 PEL
Generations ago
Relative gain = 0.551, P value = 0.024
